# Systems analyses of key metabolic modules of floral and extrafloral nectaries of cotton

**DOI:** 10.1101/857771

**Authors:** Elizabeth C. Chatt, Siti-Nabilla Mahalim, Nur-Aziatull Mohd-Fadzil, Rahul Roy, Peter M. Klinkenberg, Harry T. Horner, Marshall Hampton, Clay J. Carter, Basil J. Nikolau

**Affiliations:** Department of Biochemistry, Biophysics and Molecular Biology, Iowa State University, Ames, IA, United States; Department of Plant and Microbial Biology, University of Minnesota Twin Cities, St. Paul, MN, United States; Department of Genetics, Development and Cell Biology, Iowa State University, Ames, IA, United States; Roy J. Carver High Resolution Microscopy Facility, Iowa State University, Ames, IA, United States; Department of Mathematics & Statistics, University of Minnesota Duluth, Duluth, MN, United States

## Abstract

Nectar is a primary reward mediating plant-animal mutualisms to improve plant fitness and reproductive success. In *Gossypium hirsutum* (cotton), four distinct trichomatic nectaries develop, one floral and three extrafloral. The secreted floral and extrafloral nectars serve different purposes, with the floral nectar attracting bees to promote pollination and the extrafloral nectar attracting predatory insects as a means of indirect resistance from herbivores. Cotton therefore provides an ideal system to contrast mechanisms of nectar production and nectar composition between floral and extrafloral nectaries. Here, we report the transcriptome, ultrastructure, and metabolite spatial distribution using mass spectrometric imaging of the four cotton nectary types throughout development. Additionally, the secreted nectar metabolomes were defined and were jointly composed of 197 analytes, 60 of which were identified. Integration of theses datasets support the coordination of merocrine-based and eccrine-based models of nectar synthesis. The nectary ultrastructure supports the merocrine-based model due to the abundance of rough endoplasmic reticulum positioned parallel to the cell walls and profusion of vesicles fusing to the plasma membranes. The eccrine-based model which consist of a progression from starch synthesis to starch degradation and to sucrose biosynthesis was supported by gene expression data. This demonstrates conservation of the eccrine-based model for the first time in both trichomatic and extrafloral nectaries. Lastly, nectary gene expression data provided evidence to support *de novo* synthesis of amino acids detected in the secreted nectars.

**One sentence summary:** The eccrine-based model of nectar synthesis and secretion is conserved in both trichomatic and extrafloral nectaries determined by a system-based comparison of cotton (*Gossypium hirsutum*) nectaries.

## Introduction

Nectars are sugar-rich solutions, produced and secreted from nectary glands, and present an attractive reward to animal mutualists in exchange for ecosystem services. In the case of floral nectar this service is pollination, and extrafloral nectars are offered to recruit pugnacious predatory insects and provide indirect protection from herbivores (Mitchell et al., 2009; Ollerton, 2017; Simpson and Neff, 1981). These plant-animal mutualisms improve plant fitness and reproductive success. Domesticated Upland cotton, *Gossypium hirsutum*, develops a floral and three distinct extrafloral nectaries, all of which are trichomatic nectaries secreting the nectar from specialized papillae, a type of multicellular glandular trichome. While cotton is largely a self-pollinating crop, honey bee visitations, facilitated by the floral nectar reward, increases yield of total number of bolls and total lint mass produced (Rhodes, 2002). The extrafloral nectars provide a source of indirect protection by attracting aggressive predatory ants which ward off various herbivores (Bentley, 1977; González-Teuber et al., 2012; Rudgers et al., 2003; Rudgers and Strauss, 2004; Wäckers et al., 2001).

The patterns of nectar secretion vary among the different cotton nectaries, and are optimized for benefits, while minimizing the energetic cost of producing the nectar (Heil, 2011; Pleasants, 1983; Wäckers and Bonifay, 2004). The floral nectary actively secretes on the day of anthesis (Gilliam et al., 1981), whereas the extrafloral nectaries modulate nectar secretion based on the environmental stressor of insect herbivory (Wäckers and Bonifay, 2004). For example, the vegetative foliar nectary, located on the abaxial surface of the leaf midvein, displays low constitutive secretion, which is induced by herbivory (Wäckers et al., 2001; Wäckers and Bonifay, 2004). In contrast, the reproductive extrafloral nectaries, bracteal and circumbracteal, which are located on the abaxial surface of the bracts and sepal respectively, display peak nectar production on the day of anthesis and continue to secrete as the boll matures, but secretion will decrease in response to herbivory (Wäckers and Bonifay, 2004), indicating more complex regulatory circuitry for control in nectar production.

The molecular underpinnings of nectar synthesis and secretion are beginning to be elucidated through advancements in “omics” technologies primarily using the floral nectaries of Arabidopsis, *Cucurbita pepo* and *Nicotiana* spp.(Kram et al., 2009; Lin et al., 2014; Ren et al., 2007; Solhaug et al., 2019). These studies provide evidence to support an eccrine-based model of nectar synthesis and secretion, which utilizes pores and transporters for movement of pre-nectar metabolites through the plasma membrane of nectariferous parenchyma tissues [reviewed by (Roy et al., 2017)]. In this model, prior to nectar secretion the ‘pre-nectar’ sugar metabolites are delivered through the vasculature and stored in the nectary parenchyma, primarily as starch (Chatt et al., 2018; Lin et al., 2014; Peng et al., 2004; Ren et al., 2007; Solhaug et al., 2019). At the time of nectar secretion, the stored starch is rapidly degraded, and the products are used to synthesize sucrose through the enzymatic action of sucrose-phosphate synthases (SPS) and sucrose-phosphate phosphatases. The sucrose is exported into the apoplasm in a concentration dependent manner via the uniporter SWEET9, and subsequently hydrolyzed by cell wall invertase (CWINV4), to the hexose components glucose and fructose, thereby maintaining the sucrose concentration gradient (Lin et al., 2014; Ruhlmann et al., 2010). The last step of sucrose hydrolysis is critical to the production of hexose-rich nectars (Ruhlmann et al., 2010), but may play a minimal role in production of sucrose-rich nectars (Chatt et al., 2018; Solhaug et al., 2019).

In order to fulfill biological functions, nectar components must be released from the nectary into the environment. Nectaries containing ‘nectarostomata’ simply release the nectar through these modified stomata (Paiva, 2017). The means by which nectar passes through the cell wall and cuticle of trichomatic nectaries is unclear as the current understanding is based solely on ultrastructural analyses (Eleftheriou and Hall, 1983a; Findlay et al., 1971b; Kronestedt et al., 1986; Wergin et al., 1975). These studies indicate that at the time of nectar secretion, the cuticle separates from the cell wall on the terminal cells of the glandular trichome. Nectar then accumulates in the space between the cuticle and cell wall thereby generating hydrostatic pressure for the release of nectar as discrete droplets through the porous cuticle. It is unclear if the cell wall and cuticle undergo biochemical alterations to facilitate this process or if it is driven purely by physical force causing the cuticle to rupture.

In this study, we used a holistic approach to characterize the morphology, ultrastructure, and gene expression patterns of *G. hirsutum* floral and extrafloral nectaries as they develop from the pre-secretory to secretory to post-secretory stages. Gene expression data was also probed in the context of secreted nectar metabolomes, and to identify signatures of biochemical alterations in the cell wall and cuticle coinciding with facilitation of nectar secretion. Together these data were compared to the current eccrine-based model of nectar synthesis to assess for the first time whether this model is conserved among trichomatic and extrafloral nectaries.

## Results

Domesticated Upland cotton, *G. hirsutum* (TM-1), develops four types of nectaries, three are extrafloral and one is floral, and all consist of multicellular glandular trichomes, specifically called papillae. The three extrafloral nectary types, foliar, bracteal, and circumbracteal, are subcategorized as vegetative or reproductive. The vegetative foliar nectary is located on the abaxial surface of the leaf midrib (Fig. 1A; Fig. 2A, B). The bracteal and circumbracteal nectaries are reproductive extrafloral nectaries due to their close association with the flower. The bracteal nectaries, also referred to as the outer involucellar or subbracteal, develop at the base of each bract subtending the flower and framing the cotton boll (Fig. 1B; Fig. 2C, D). The circumbracteal or inner involucellar nectary occurs on the abaxial calyx surface alternate with the bracts (Fig. 1C; Fig. 2E, F). The floral nectary develops on the adaxial calyx surface and lines the basal circumference. The secretory papillae of the floral nectary subtend a ring of stellate trichomes (Fig. 2G, H).

**FIGURE 1.**
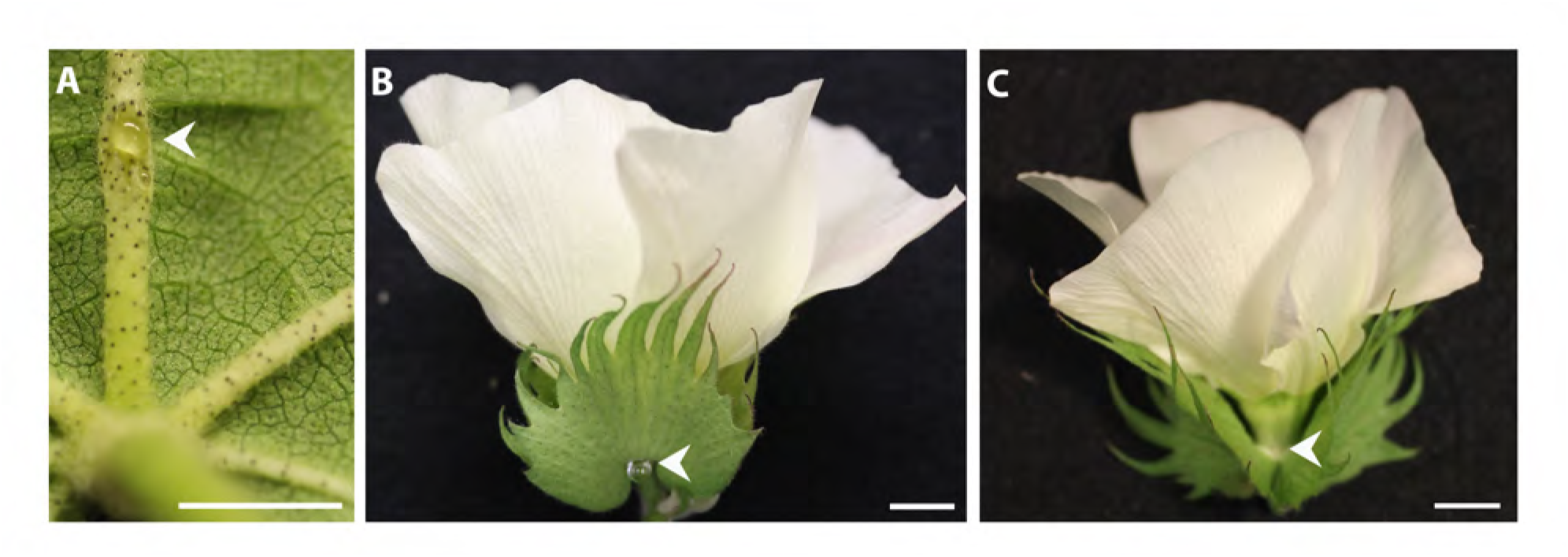
Extrafloral nectaries present on *G. hirsutum* leaves and flowers indicated by arrow heads: the foliar nectary (**A**), bracteal nectary (**B**), and circumbracteal nectary (**C**). Scale bar A = 5 mm; B, C = 10 mm.

**FIGURE 2.**
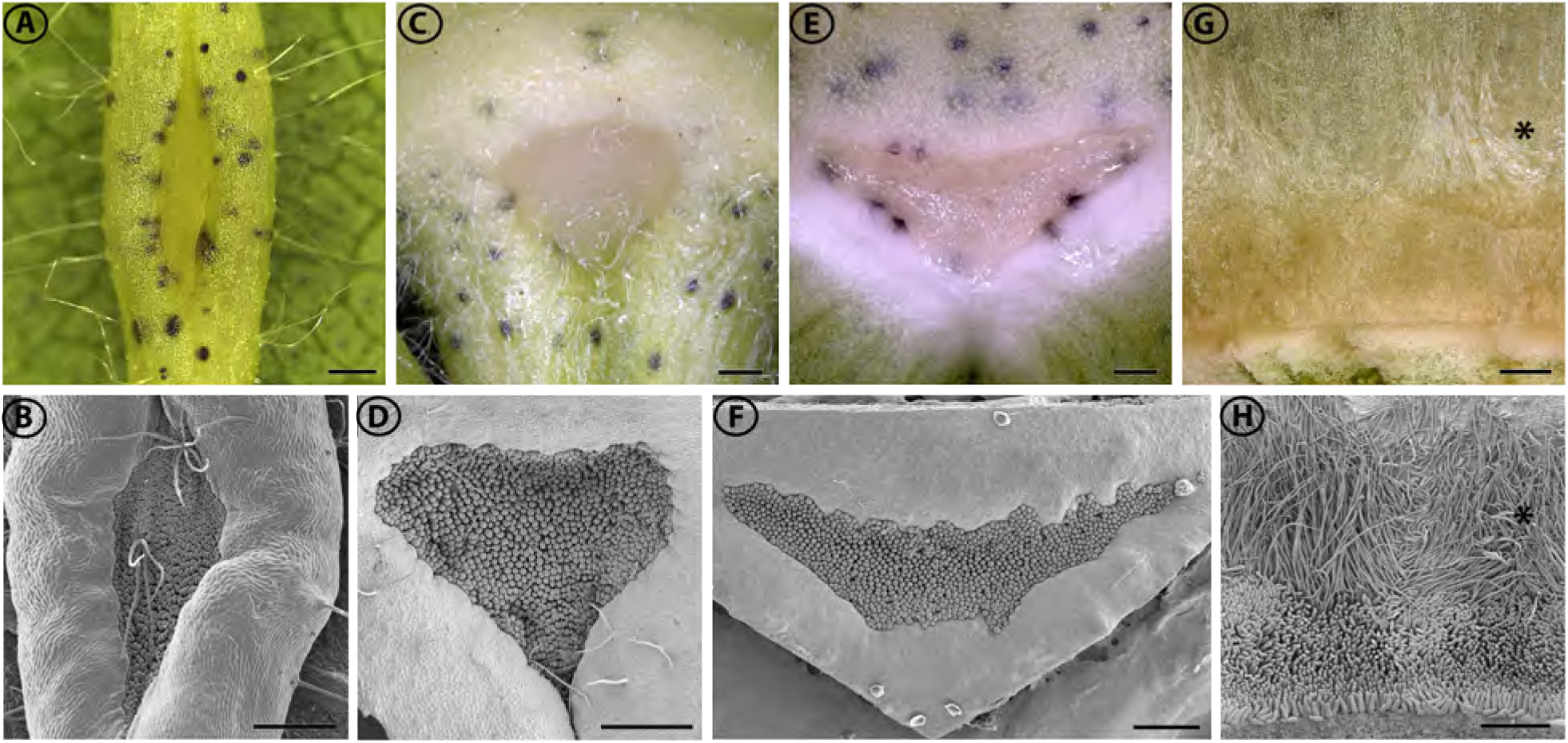
Macrostructure of *G. hirsutum* nectaries at the secretory stage of development, viewed with a macrozoom microscope (A, C, E, G) and SEM (B, D, F, H). Extrafloral nectaries (**A-F**) are composed of a pit of densely packed papillae. Floral nectary (**G, H**) is composed of a ring of stellate trichomes (*) subtended by a ring of papillae. (**A, B**) Foliar nectary; (**C, D**) Bracteal nectary; (**E, F**) Circumbracteal nectary; (**G, H**) Floral nectary. Scale bars = 0.5 mm

### General features of nectary epidermis and parenchyma

The epidermis and parenchyma of all four nectary types were examined and compared at two developmental stages, pre-secretory and secretory. The nectary epidermal tissue contains two distinct regions, one bordering the papillae and the second directly below the papillae. Epidermal tissue bordering the papillae of each nectary are highly vacuolated (Fig. 3D, E). In the floral nectaries and pre-secretory foliar nectaries, the boardering epidermal tissue contain bodies that stain heavily with Toluidine Blue and osmium tetroxide indicating the presecence of phenolics. These densely-staining bodies are 21 ± 8 µm in diameter. In contrast, the bracteal and circumbracteal nectaries lack the densely-staining bodies within the boardering epidermal tissue (Fig. 3A). The second region of epidermal tissue, the hypoepidermis located below the papillae, is characterized by densely-staining cytoplasm and the densely-staining bodies (Fig. 4B, D, F and H). At the pre-secretory stage, the extrafloral nectary hypoepidermis is vacuolated (Fig 4A, C, E), while the floral nectary hypoepidermis is not vacuolated and instead contains a dense-staining cytoplasm (4G).

**FIGURE 3.**
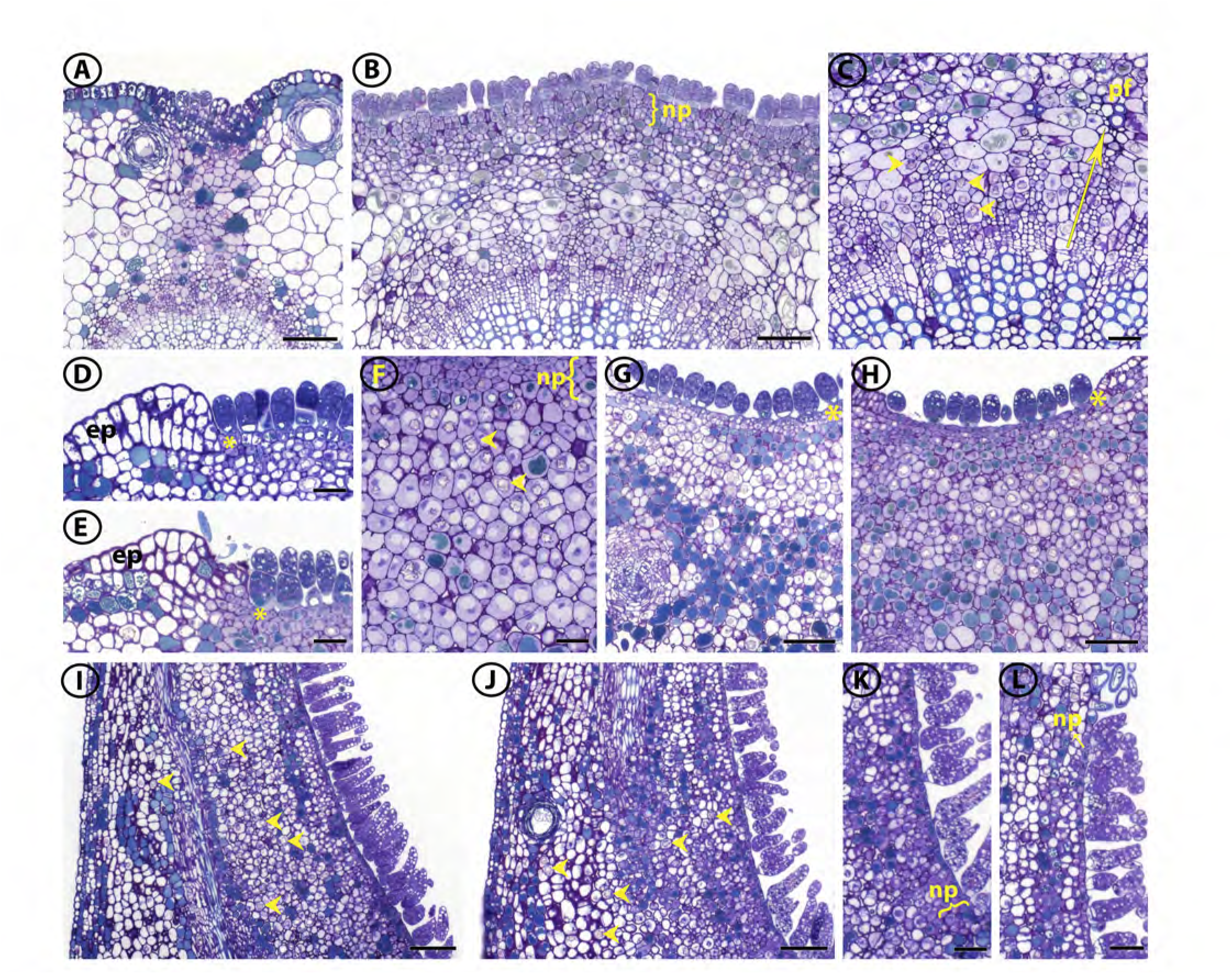
Light micrographs of *G. hirsutum* nectary longitudinal sections stained with Toluidine Blue O. (**A**) Foliar pre-secretory nectary; (**B**) Foliar secretory nectary; (**C**) Foliar secretory nectary, phloem rays extending into the subnectariferous parenchyma highlighted by arrow, arrow heads point to druse crystals; (**D**) Bracteal pre-secretory nectary; (**E**) Bracteal secretory nectary; (**F**) Bracteal secretory nectary nectariferous and subnectariferous parenchyma subtending the papillae, arrow heads point to druse crystals; (**G**) Circumbracteal pre-secretory nectary; (**H**) Circumbracteal secretory nectary; (**I**) Floral pre-secretory nectary, arrow heads point to druse crystals; (**J**) Floral secretory nectary, arrow heads point to druse crystals; (**K**) Proximal portion of floral secretory nectary; (**L**) Distal portion of floral secretory nectary. Abbreviations: ep = epidermis; np = nectariferous parenchyma; pf = phloem fiber; * = hypoepidermis. Scale bars A, B, G, H, I, J = 100 µm; C, D, E, F, K, L = 50 µm.

**FIGURE 4.**
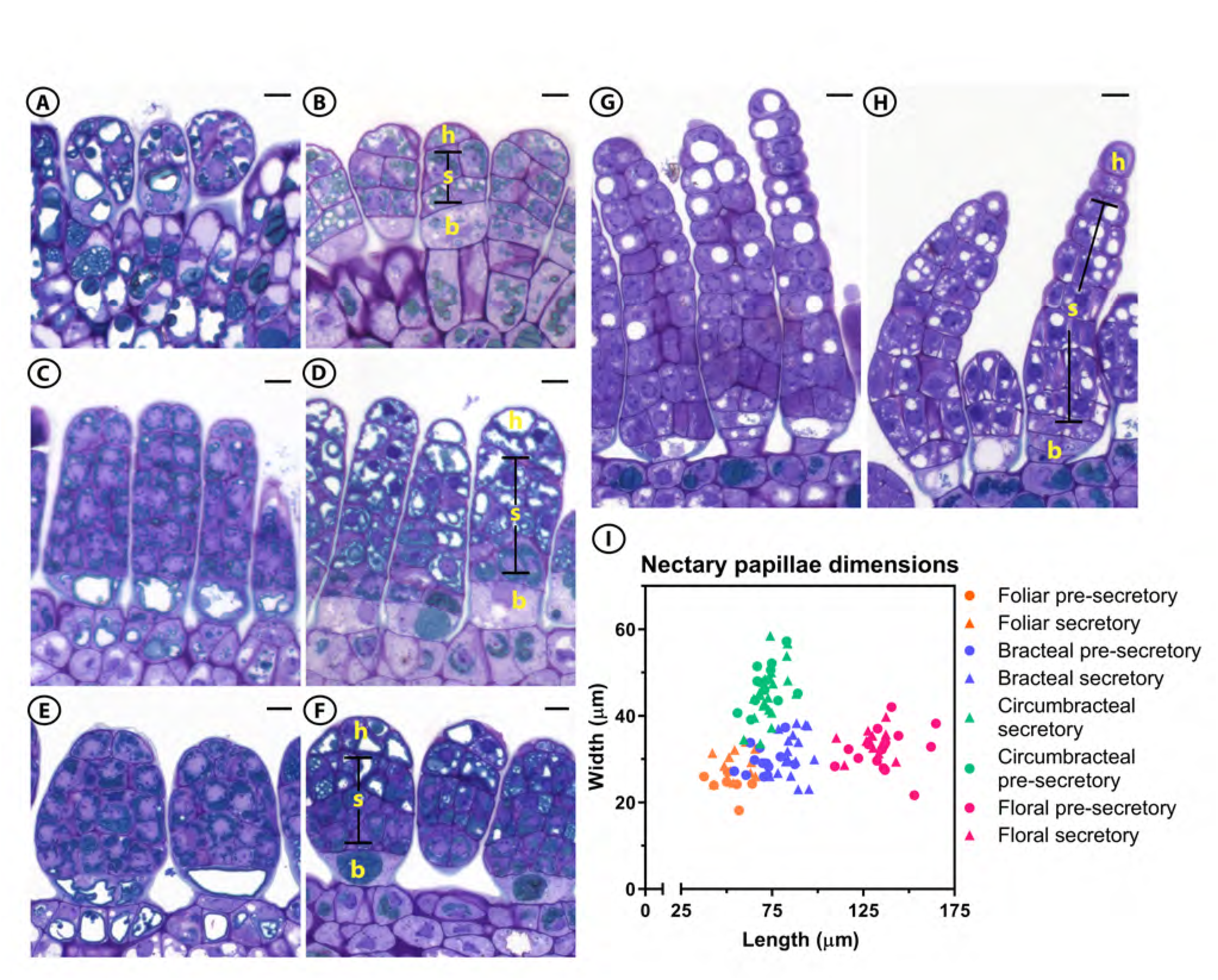
Light micrographs of *G. hirsutum* papillae longitudinal sections from the four different nectary types stained with Toluidine Blue O and their dimensions. (**A**) Foliar pre-secretory; (**B**) Foliar secretory; (**C**) Bracteal pre-secretory; (**D**) Bracteal secretory; (**E**) Circumbracteal pre-secretory; (**F**) Circumbracteal secretory; (**G**) Floral pre-secretory; (**H**) Floral secretory; (**I**) Length and width distribution of the nectary papillae at different stages of development. A total of 7 to 22 papillae were measured for each nectary type and at each developmental stage. Abbreviations: h = head cells; s = stalk cells; b = basal cells. All scale bars = 10 µm

The nectariferous parenchyma of all nectary types, located between the subnectariferous parenchyma and the secretory papillae, is characterized by isodiametric cells with minimal intercellular spaces, and containing diminutive phenolic bodies, and densely-staining cytoplasm. The number of nectariferous parenchyma layers vary among the nectary type and developmental stage. The foliar, bracteal, and circumbracteal nectaries, at the pre-secretory stage contain three to four layers of nectariferous parenchyma (Fig. 3A, D, G). This number of cell-layers is maintained at the secretory stage in the bracteal and circumbracteal nectaries (Fig. 3E, H) but increases up to six layers in the foliar nectary (Fig. 3B). The number of nectariferous parenchyma cell-layers of the floral nectary varies depending on the position within the nectary. At both developmental stages, the proximal end contains three to four cell-layers (Fig. 3K) and the number of cell-layers decreases to two at the far distal end (Fig. 3L).

The subnectariferous parenchyma is composed of approximately ten layers of large cells with intermediate cytoplasm density as compared to the nectariferous parenchyma. Vascular bundles are present near the subnectariferous parenchyma with phloem rays extending into the subnectariferous parenchyma of the foliar nectary exclusively (Fig. 3C). The subnectariferous parenchyma of all examined nectary types develop densely-staining bodies, which occur more abundantly in cells surrounding the vascular bundles (Fig. 5).

**FIGURE 5.**
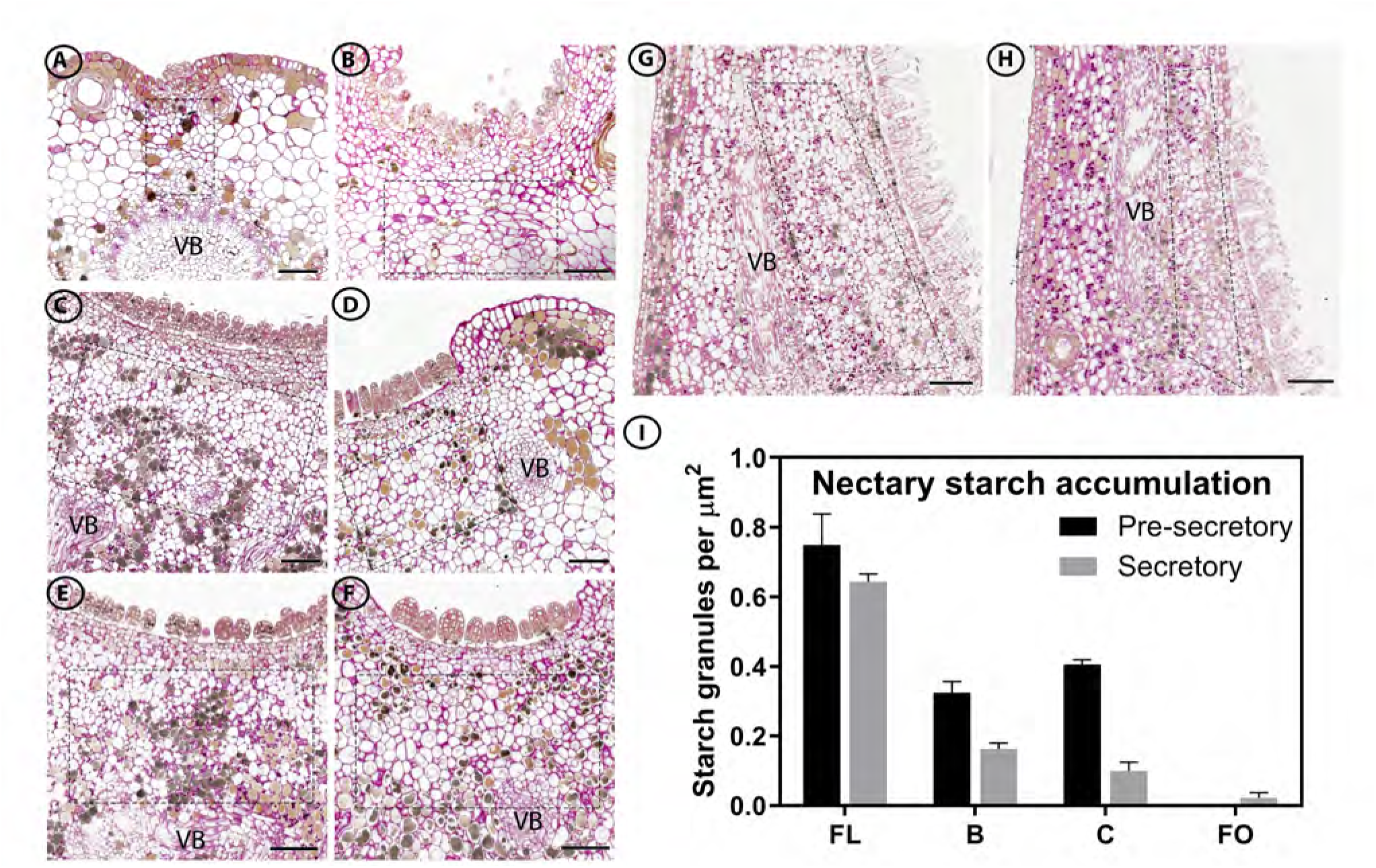
Distribution of starch granules within subnectariferous parenchyma (regions within dashed boxes) during development of *G. hirsutum* nectaries visualized by PAS staining and light microscopy and their density. (**A**) Foliar pre-secretory; (**B**) Foliar secretory; (**C**) Bracteal pre-secretory; (**D**) Bracteal secretory; (**E**) Circumbracteal pre-secretory; (**F**) Circumbracteal secretory (**G**) Floral pre-secretory; (**H**) Floral secretory; (**I**) Density of starch granules within the subnectariferous parenchyma of *G. hirsutum* nectaries during nectary development. For each nectary type and developmental stage, starch granules were counted from a minimum of six sections originating from two separate nectaries. Error bars represent S.E. Abbreviations: VB = vascular bundle; FL = floral; B = bracteal; C = circumbracteal; FO = foliar. Scale bars = 100 µm.

Cells containing druses (spherical aggregates of calcium oxalate crystals) were primarily observed in the subnectariferous parenchyma of all nectary types, especially surrounding the vascular bundles. The druses present in foliar nectaries align in a row, in parallel to the phloem rays from the vascular bundles to the papillae (Fig. 3C). Druses were most abundant in the floral nectaries, occurring throughout the subnectariferous and nectariferous parenchyma (Fig. 3I, J).

Starch accumulation within the nectaries was visualized by PAS staining. Starch granules occur in the subnectariferous parenchyma of reproductive nectaries, floral, bracteal, and circumbracteal. These are most commonly located near the vascular bundles of the bracteal and circumbracteal nectaries (Fig. 5C, E), and the frequency of these granules decrease as the nectaries transition from the pre-secretory to the secretory stages (Fig. 5D, F, I). Floral nectaries accumulate larger starch granules in the subnectariferous parenchyma at both stages of development with a slight decrease at the secretory stage (Fig. 5G, H, I). In contrast, virtually no starch granules were observed within the subnectariferous parenchyma of the foliar nectaries at either developmental stages (Fig. 5A, B, I).

### Morphological features of nectary papillae

The papillae of all nectaries are multicellular and contain three regions typical of glandular trichomes; these being basal cell(s), stalk cells, and head cells. Mature extrafloral papillae contain five to six layers of cells with an average papillae-length of 68 ± 14 µm (SD), while the floral papillae are more extensive, with 12 to 14 cell layers with an average papillae-length of 133 ± 10 µm (SD) (Fig. 4B, D, E, H).

Regardless of papillae-length, each papilla begins with distinct basal cell(s), which lack electron-dense cytoplasm. The three types of extrafloral nectaries contain a single basal cell (Fig. 4A-F), while the floral nectary contains two basal cells (Fig. 4G,H; Fig. 7G). The stalk cells, characterized by phenolic bodies and vacuoles, determine the papillae length and width, and the circumbracteal nectaries have the widest papillae (46 ± 6 µm), as compared to the papillae of the other three nectaries (30 ± 4 µm) (Fig. 4I). The densely-staining bodies in the stalk cells of the bracteal and circumbracteal nectaries are arranged around the cell periphery (Fig. 4C-F).

**FIGURE 7.**
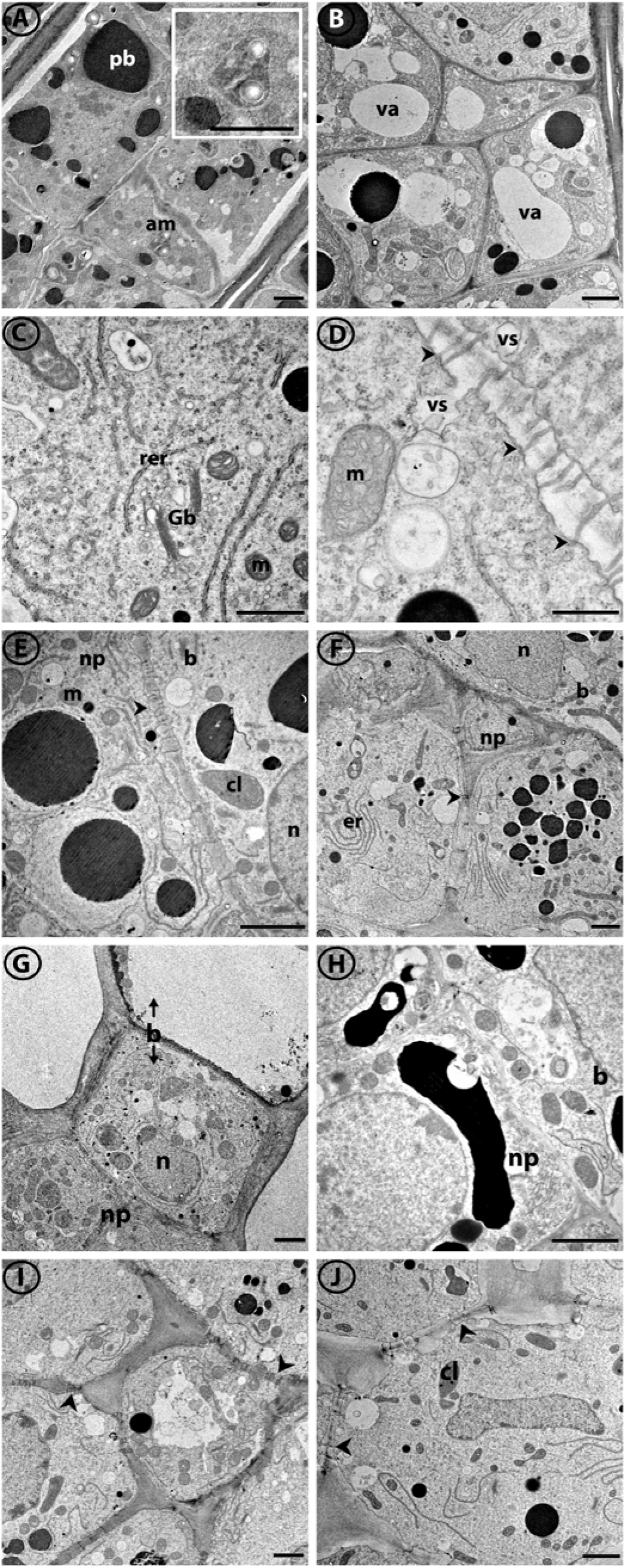
TEM of the cellular details of *G. hirsutum* nectary papillae and supporting nectariferous parenchyma tissue at the secretory stage. (**A**) Stalk cells from bracteal nectary with amyloplast insert; (**B**) Stalk cell from foliar nectary; (**C**) Organelles of stalk cell exemplified by foliar nectary; (**D**) Plasmodesmata (arrowheads) in cell wall of internal stalk cell; (**E-G**) Junction between basal cell and nectariferous parenchyma of (**E**) bracteal nectary; (**F**) foliar nectary; (**G**) floral nectary; (**H**) Basal cell from circumbracteal nectary; (**I**) Nectariferous parenchyma from bracteal nectary; (**J**) Nectariferous parenchyma from foliar nectary. Arrowheads identify plasmodesmata. Abbreviations: am = amyloplasts; cl = chloroplast; b = basal cell; er = endoplasmic reticulum; Gb = Golgi body; m = mitochondria; n = nucleus; np = nectariferous parenchyma; pb = phenolic body; rer = rough endoplasmic reticulum; va = vacuole; vs = vesicle. Scale bars A, B, E-J = 2 µm; C = 1 µm; D = 0.5 µm.

The size and number of vacuoles differ among the different types of nectaries and their stages of development. Pre-secretory stalk and head cells of bracteal and circumbracteal nectaries contain virtually no vacuoles (Fig. 4C, E), while at the secretory stage the distal two-thirds of the papillae cells become highly vacuolated, especially the head cells (Fig. 4D, H). In contrast to the bracteal and circumbracteal nectaries, the pre-secretory stalk and head cells of foliar and floral nectaries contain large, circular vacuoles in section (Fig. 4A, G), and by the secretory stage these vacuoles become smaller, and more numerous within the cells of the distal two-thirds of the papillae (Fig. 4B, H).

The cuticle and cell wall of the papillae have notable characteristics that are common among the four nectaries. These cuticles are thinnest around the head cell (Fig. 6G) and become thicker at the basal cell-epidermis junctions (Fig. 7G). Furthermore, in all four nectary types, at the secretory stage the cuticle of the head cells separates from the underlying cell wall and displays microchannels (Fig. 6). These microchannels are visible as slits on the outer surface of the papillae head cells (Fig. 6B, C). In the case of the bracteal and circumbracteal nectaries this separation of the cuticle occurs earlier in development, at the pre-secretory stage, and occasionally extends down to the distal stalk cells of bracteal papillae. Cell wall ingrowths toward the plasma membrane were observed in the bracteal and circumbracteal papillae head cells at the secretory stage (Fig. 6D, E). Occasionally an extensive periplasmic space is present in the bracteal stalk cells (Fig. 6F).

**FIGURE 6.**
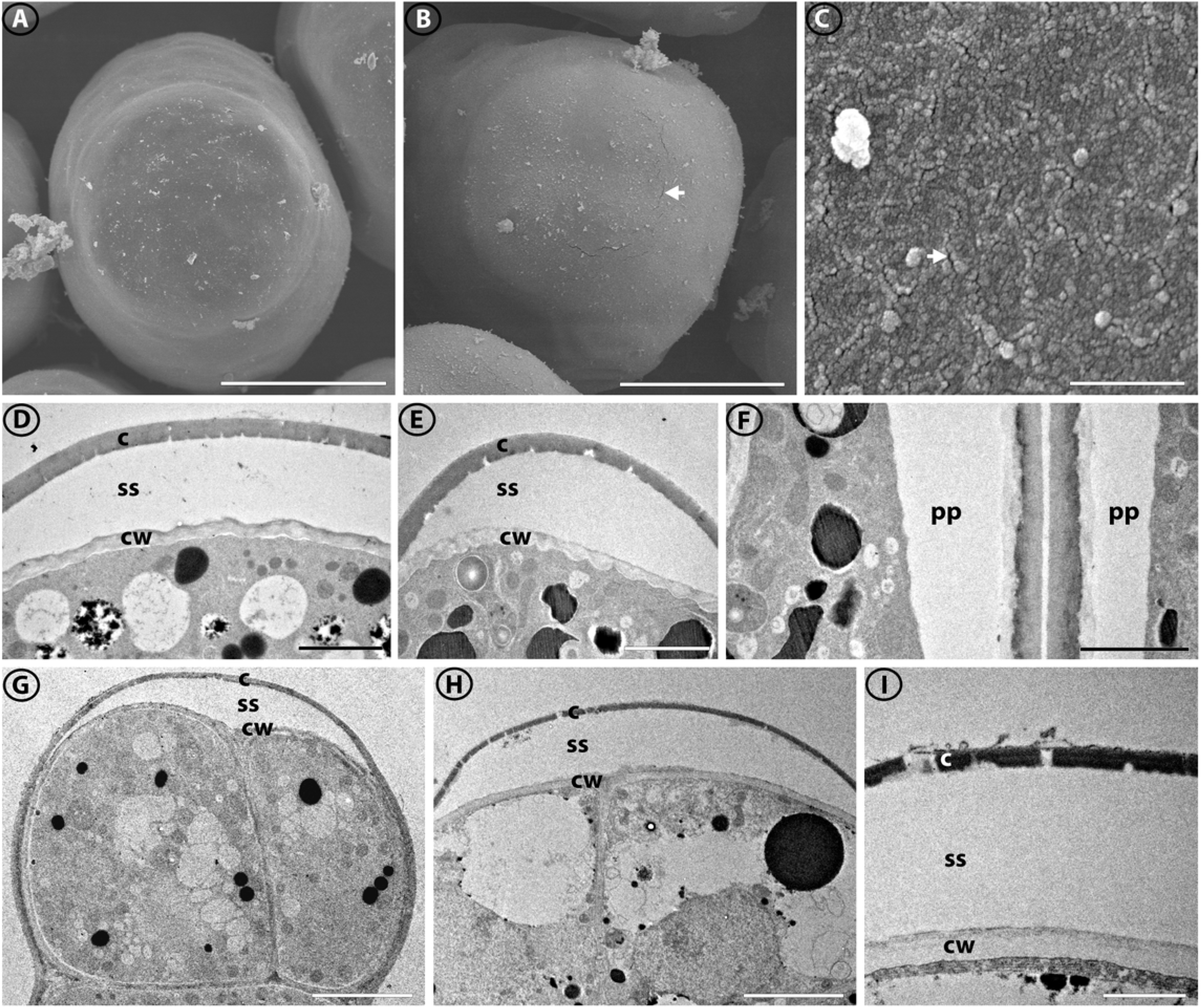
SEM (A-C) and TEM (D-I) images of the cuticle and cell wall of *G. hirsutum* nectary papillae. (**A**) Terminal end of papillae of circumbracteal nectary at pre-secretory stage, note lack of microchannels (cracks) in cuticle surface; (**B**) Circumbracteal nectary at secretory stage, arrow head identifies the microchannels in the cuticle surface; (**C**) Surface of a terminal cell from a foliar nectary papilla at secretory stage, arrowhead identifies the cuticular microchannels; (**D**) Head cell from secretory circumbracteal papilla; (**E**) Secretory bracteal papilla showing separated cuticle (c) with microchannels and cell wall ingrowths; (**F**) Two adjacent distal stalk cells from secretory bracteal papillae, note periplasmic space (pp); (**G**) Head cells from secretory floral papilla; (**H**) Secretory foliar papilla showing separated cuticle; (**I**) Porous cuticle of head cell of bracteal secretory papilla. Abbreviations: c = cuticle; cw = cell wall; ss = subcuticular space; pp = periplasmic space. Scale bars A, B = 25 µm; C = 50 nm; D, E, F = 2 µm; G, H = 5 µm; I = 1 µm

### Organelle composition

The organelle composition of the papillae glands and supporting nectariferous parenchyma was examined by TEM. Cells of the papillae from all cotton nectaries are nucleated. The most common organelles observed in these cells are mitochondria, rough endoplasmic reticulum, and vesicles (Fig. 7C, D), whereas Golgi bodies (Fig. 7C) and amyloplasts (Fig. 7A) are significantly less abundant, and simple chloroplasts only occur in the bracteal (Fig. 7E) and foliar (Fig. 7J) nectaries. Among the four types of nectaries, the cells of the floral nectary appears to have the most mitochondria, and among all the nectaries, the mitochondria are typically located around the cell periphery in close proximity to rough endoplasmic reticulum. The basal cells of the papillae glands appear to have higher organelle complexity, containing more mitochondria and rough endoplasmic reticulum per cell (Fig. 7E-H), while the head cells display the least organelle complexity (Fig. 6D, E, G, H). Throughout the papillae and nectariferous parenchyma, vesicle fusion to the plasma membrane was frequently observed (Fig. 7D), and typical of nectary tissue, plasmodesmata traverse the inner anticlinal and peridermal walls of these tissues (Fig. 7D, E, I, J).

### Nectar metabolome

Metabolomics analysis of the nectar from the four cotton types led to the detection and quantification of 197 analytes, with the successful chemical identification of 60 metabolites (Supplemental File 1). These latter metabolites include the dominant sugars, and the minor components, which are amino acids, sugar alcohols, lipids, diols, organic acids, esters, and aromatics.

The major constituents of the four nectars are similar, being hexose-dominant, with an equal molar ratio of fructose:glucose (Table 1). However, the four different nectar types can be distinguished based on the minor nectar metabolites, particularly between the floral and the extrafloral nectars (Supplemental Fig. 1 and 2). Variation between the three nectar categories is clearly illustrated by sucrose abundance, which differs significantly between floral, reproductive extrafloral, and vegetative extrafloral nectars (Table 1). These compositional variations are visualized by the pairwise volcano plots shown in Figure 8, which reveal that 105 of the 197 detected analytes significantly differ in abundance in at least one pairwise comparison (q-value < 0.05, Supplemental File 2). The floral nectar is compositionally distinct from the extrafloral nectars, with at least 77 distinguishing analytes between each extrafloral nectar from the floral nectar (Fig. 8A-C). Specifically, the amino acids are more abundant in the floral nectar (Table 1; Supplemental Fig. 2, Cluster 8), particularly aspartic acid, asparagine, leucine, phenylalanine, tryptophan, and gamma-aminobutyric acid (GABA) occurring exclusively in the floral nectar (Supplemental Fig. 1; Supplemental File 2). The other distinguishing compositional difference among these amino acids is the finding that the extrafloral nectars are less abundant in non-proteinaceous and essential amino acids (Supplemental Fig. 3).

**FIGURE 8.**
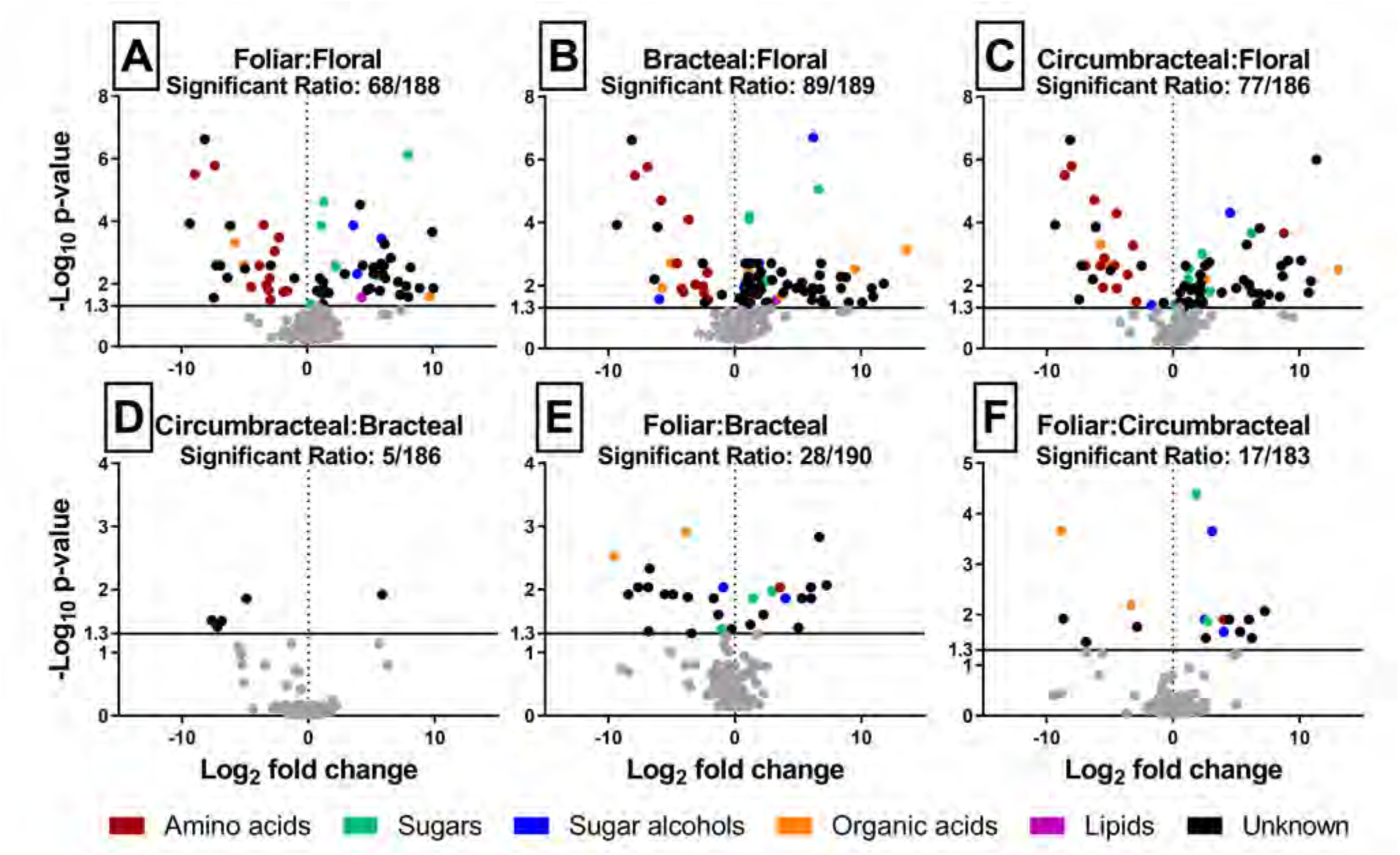
Volcano plot analyses of all possible pairwise comparisons of *G. hirsutum* nectar metabolomes. In each comparison, “significant ratio” identifies the proportion of the detected analytes whose abundance difference is statistically significant (colored data points above the y-axis value of 1.3) between the two nectar types. The chemical class identity of the metabolites is color-coded.

**Table 1.** Predominant sugars, and amino acids in *G. hirsutum* nectars. Different superscript letters indicate statistically significant differences in abundance (q-value < 0.05).

| Nectar Type | Sugars (M) |  |  | Fructose-to-glucose ratio | Sucrose-to-hexose ratio | Amino acids (μM) |  |  |  |
| --- | --- | --- | --- | --- | --- | --- | --- | --- | --- |
|  | Fructose | Glucose | Sucrose |  |  | Essential | Non-essential | Non-proteinaceous | Total |
| Floral | 1.81 ± 0.14<sup>A</sup> | 1.89 ± 0.18<sup>A</sup> | 0.005 ± 0.001<sup>A</sup> | 0.97 ± 0.03 | 0.0014 ± 0.0004 | 116 ± 11 | 2950 ± 294 | 3.9 ± 1.2 | 3070 ± 303 |
| Bracteal | 4.05 ± 0.32<sup>B</sup> | 4.27 ± 0.32<sup>B</sup> | 0.50 ± 0.06<sup>B</sup> | 0.95 ± 0.01 | 0.060 ± 0.005 | 12 ± 7 | 37 ± 12 | 5.7 ± 0.9 | 54 ± 20 |
| Circumbracteal | 4.3 ± 0.6<sup>B</sup> | 4.3 ± 0.7<sup>B</sup> | 0.37 ± 0.07<sup>B</sup> | 1.02 ± 0.03 | 0.040 ± 0.004 | 1.9 ± 0.4 | 21 ± 3 | 6.8 ± 2.8 | 30 ± 2 |
| Foliar | 4.5 ± 0.4<sup>B</sup> | 4.2 ± 0.3<sup>B</sup> | 1.3 ± 0.1<sup>C</sup> | 1.10 ± 0.01 | 0.150 ± 0.003 | 11 ± 6 | 40 ± 13 | 3 ± 1 | 55 ± 15 |

### Mass spectrometric imaging of nectary metabolite distribution

The application of MALDI-based mass spectrometric imaging technology on the four nectary types at two developmental stages (pre-secretory and secretory), resulted in the detection of over 7,000 ion-analytes, each of which are distinguishable by their unique *m/z* values. This dataset was refined by applying two selection filters in order to reduce the number of ion-analytes and begin the process of idetifying the chemical nature of each ion. One of these filters evaluated the “reliability” of ion-detection from nectar tissue associated pixels. Namely, ions that were detectable in 5 out of 10 near-adjoining pixels, which were positioned over papillae gland or nectariferous parenchyma nectar tissues were were considered reliable and were retained. The second filter compared the ion-strengths of each ion from tissue associated pixels to the signal strength obtained from non-tissue pixels, retaining only those ions that showed 2-times greater signal strength from tissue-pixels compared to background signal obtained from pixels devoid of tissue. Implementing these criteria reduced the dataset to 161 analytes of distinct *m/z* values. The distribution of these 161 analytes across the nectary tissues is not uniform, indicating the heterogeneity in the metabolic status of the cells within each nectary (Supplemental Fig. 4). The chemical nature of 101 of these ions were tentatively identified (Supplemental File 3) based on the accurate mass of each ion (Δppm ≤ 8), as compared to entries in the METLIN chemical database (https://metlin.scripps.edu). Approximately 60% of these tentatively identified analytes are phenolic type metabolites, and they are localized near the vasculature within the subnectariferous parenchyma and the epidermis (Fig. 9). This distribution matches the distribution of subcellular bodies that stain heavily with osmium tetroxide and are visualized by TEM (Fig. 7), confirming their identity as polyphenolic compounds.

**FIGURE 9.**
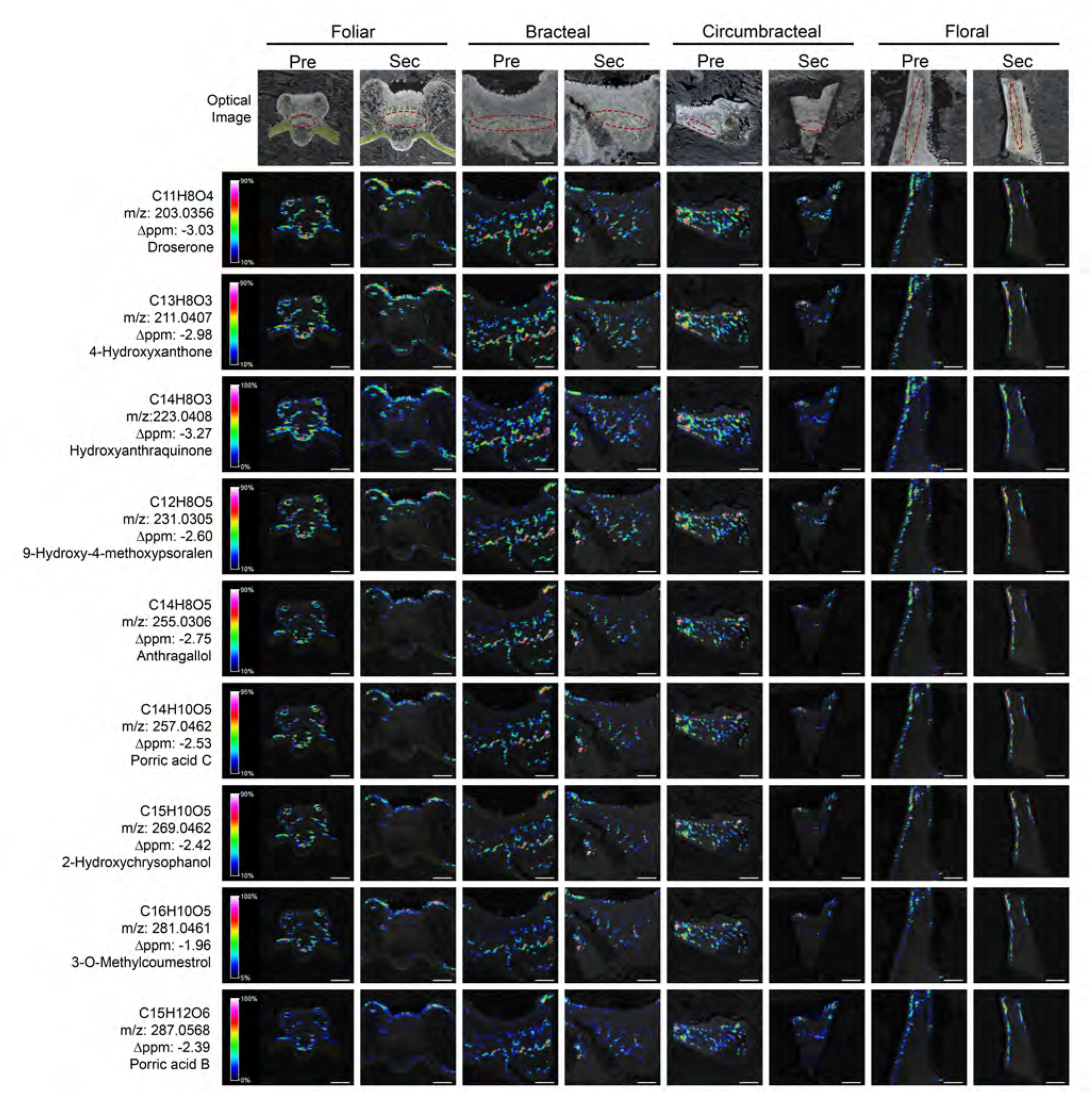
Spatial distribution of phenolic metabolites visualized by mass-spectrometric imaging. Each MS image was obtained from the longitudinal cryosections of *G. hirsutum* nectaries that were optically imaged in parallel (top row). The position of the vasculature is highlighted by red colored ovals in the optical images. The MS imaging data was collected with a laser spot size, enabling a 25-µm spatial resolution of the metabolites. The ion signals are scaled to the maximum signal of the highest spectrum. The scaled ion signals are displayed by the rainbow heat map coloration. Scale bars = 500 µm.

### RNA sequencing and differential expression analyses

The transcriptomes of the four cotton nectary types were resolved through three development stages using RNA-seq. Over 360M sequencing reads (125 bp, paired end) were generated from RNA isolated from the four cotton nectaries and from the adjacent non-nectary control tissue. These reads were initially mapped to the UTX-JGI *G. hirsutum* genome (v1.1) and sunsequently mapped to *Arabidopsis thaliana* Col-0 genome. The latter was selected because the Arabidopsis genome is well annotated and has served as the genetic model for plant biology, including the process of nectar production [reviewed in (Roy et al., 2017)] (Supplemental File 4).

The DESeq statistical package (Anders and Huber, 2010) was used to identify differentially expressed genes between each nectary type and the adjacent non-nectary control tissue, and these were also compared to evaluate the effect of development on each nectary type (Supplemental File 5 and 6). These analyses revealed genes that are differentially expressed among the four nectaries at each specific stage of maturation, and those that are commonly nectary-enriched, irrespective of the four nectary types (Fig. 10).

**FIGURE 10.**
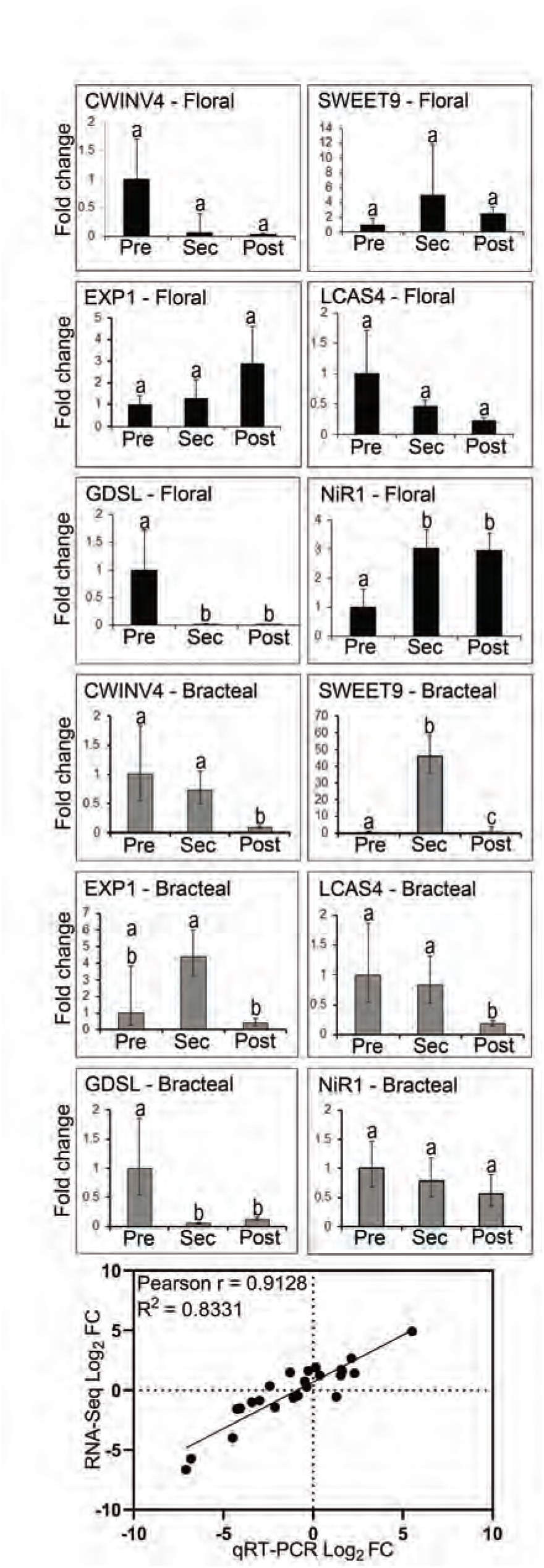
Validation of RNA-seq data by parallel qRT-PCR analysis. Using the identical RNA samples subjected to RNA-seq analysis, the expression of 6 targeted genes was analyzed by qRT-PCR. These genes are: *CWINV4* (*Cell Wall Invertase 4*), *EXP1* (*Expansin1*); *NiR1* (*Nitrite reductase 1*); *LCAS4* (*long chain acyl-CoA synthetase 4-like*); *GDSL* (*GDSL-like* Lipase/acylhydrolase); and *SWEET9* (*Sugars Will Eventually be Exported Transporter 9*). Expression was evaluated during the development of floral and bracteal nectaries as they transition from pre-secretary (Pre) to secretory (Sec) and to post-secretary (Post) stages, and the data are expressed as fold-change relative to the pre-secretory stage. Error bars represent SE from a total of 3 biological replicates. The scatter plot displays the Pearson’s correlation analysis between the RNA-seq and qRT-PCR datasets, expressed as fold-change in expression relative to the pre-secretory stage (on a log base-2 scale).

Expression profiles identified via RNA-seq analysis were validated by quantitative real time PCR (qRT-PCR) analysis using RNA isolated from floral and bracteal nectaries. These validation genes were chosen based on their known or suspected functionality in nectary development (Kram et al., 2008; Lin et al., 2014; Ruhlmann et al., 2010; Solhaug et al., 2019). Some of the selected genes display distinctive differential expression during nectary development, while others show a more stable expression pattern (e.g., *NiR1*). The qRT-PCR expression data for these six selected genes were compared to the RNA-Seq expression values obtained from the floral and bracteal nectaries from different developmental stages. Pearson’s correlation analysis of these two datasets leads to the finding of a strong positive correlation between these two methods of measuring gene expression (R^2^ = 0.83; Fig. 10). Therefore these validations indicate that the RNA-seq analyses can be used to draw conclusions concerning gene expression activity in developing nectary tissues.

A total of 3,340 genes displayed differential expression patterns between the nectary tissue and the adjacent non-nectary control tissue for at least one pairwise comparison (grey data-points in Figure 11A). These genes however, did not demonstrate any temporal change in expression during the development of each nectary-type. A summary of these genes and their occurrence among the four nectary types is visualized as a Venn diagram in Fig. 11B (Supplemental File 7). Gene ontology analysis of these genes that are commonly differentially expressed between nectary and non-nectary tissues among all four nectary types reveals an enrichment for molecular functions and biological process terminologies related to oxidoreductase activities, which are consistent with the need to generate nectar precursor metabolites and cellular energetics (Supplemental File 8). We surmise therefore these are basal functionalities that are commonly required in maintaining an operational nectary.

**FIGURE 11.**
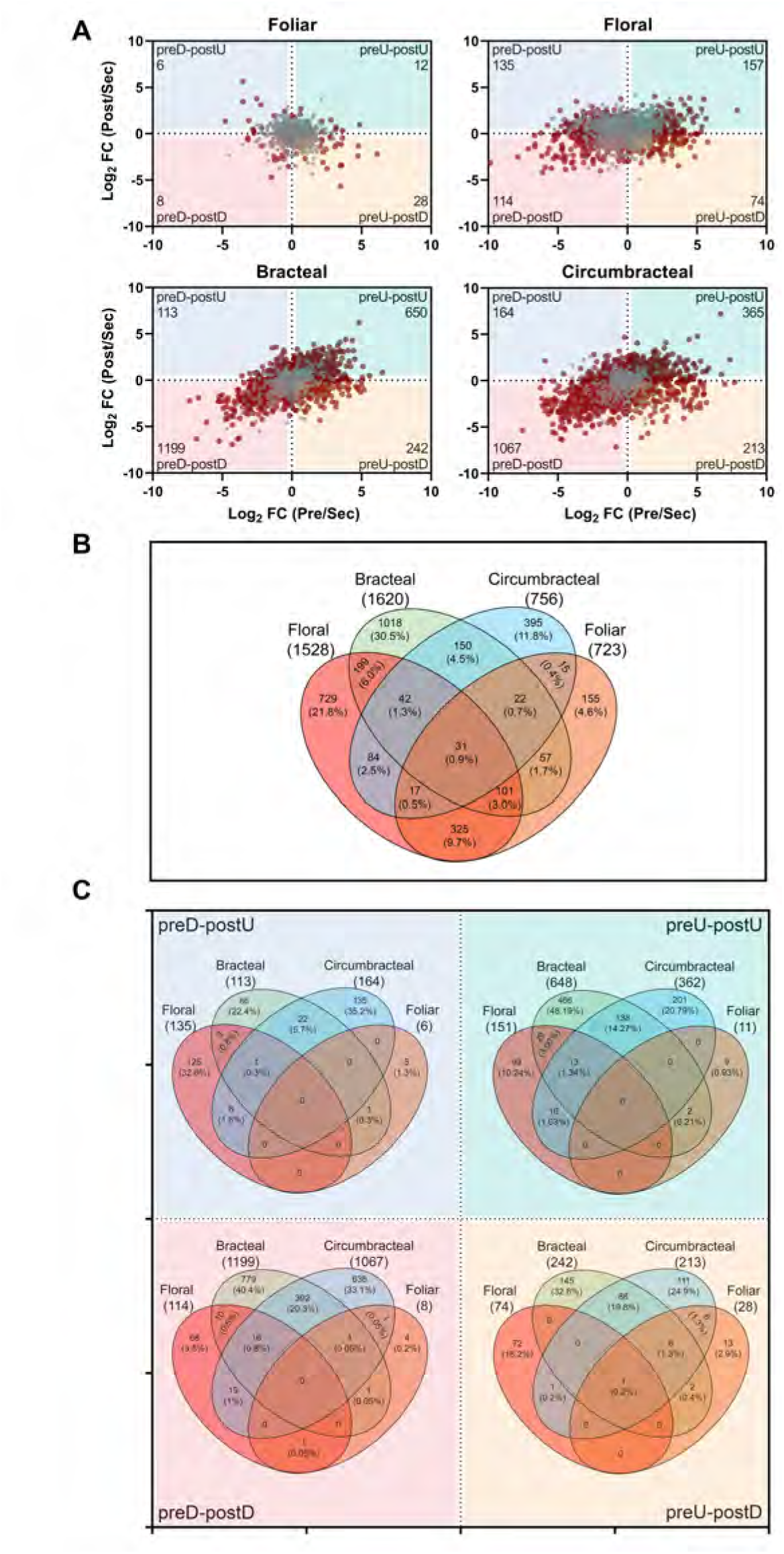
Differentially expressed genes in four nectary types. (**A**) Scatter plots displaying differentially expressed genes in relation to the development of each nectary from pre-secretory (Pre) to secretory (Sec) to post-secretory (Post) stages, normalized to the expression level at the secretory stage. Grey colored data points represent genes that are preferentially expressed in each nectary type with respect to the adjoining non-nectary tissue, but expression is minimally affected by nectary development. Red colored data points represent genes that are differentially expressed in each nectary type, and expression is also modulated by the development of each nectary type. These red data points are divided into four quadrants, which detail changes in gene expression patterns normalized to the secretory developmental stage:1) down-regulated at the pre-secretory stage and up-regulated at the post-secretory stage (preD-postU); 2) up-regulated at the pre-secretory stage and up-regulated at the post-secretory stage (preU-postU); 3) up-regulated at the pre-secretory stage and down-regulated at the post-secretory stage (preU-postD); and 4) down-regulated at the pre-secretory stage and down-regulated at the post-secretory stage (preD-postD). The number of differentially expressed genes in each sector is identified in the outer corner of each sector. (**B**)Venn diagram representation of the distribution of genes displaying nectary tissue preferential expression, but not modulated by the developmental stage of each nectary (i.e., the genes identified by grey data-points in panel A). The digits identify the absolute number and percentage of genes falling into each subset category. (**C**) Venn diagram representation of the distribution of genes that show nectary-tissue specific expression and temporal patterns of gene expression as they transition through presecretory, secretory and post-secretory stages of development (i.e., overlap among the genes represented by red-colored data-points in panel A.) The digits identify the absolute number and percentage of genes falling into each subset category.

The numbers of genes displaying a temporal change in expression, from pre-secretory to secretory to post-secretory stages associated with each nectary type are identified as red data-points in the scatter plots shown in Figure 11A (Supplemental File 6). Each scatter plot is divided into quadrants detailing the following four temporal patterns of gene expression relative to the secretory stage: 1) down-regulated at the pre-secretory stage and up-regulated at the post-secretory stage (preD-postU); 2) up-regulated at the pre-secretory stage and up-regulated at the post-secretory stage (preU-postU); 3) up-regulated at the pre-secretory stage and down-regulated at the post-secretory stage (preU-postD); and 4) down-regulated at the pre-secretory stage and down-regulated at the post-secretory stage (preD-postD). The Venn diagrams in Figure 11C (Supplemental File 9) identify the number of genes that share common temporal patterns of gene expression among the four nectary types.

These comparisons indicate that each nectary type displays a distinct temporal program of gene expression as they develop from pre-secretory to post-secretory stages. For example, there is only a single gene, terpene synthase 21 (AT5G23960.2), which shares the same temporal expression pattern across all four nectary types. Analogously, the bracteal and circumbracteal nectaries display temporal gene expression profiles that are most similar to each other (sharing 17% of the differentially expressed genes), while the floral and vegetative foliar nectaries are most distinct (sharing only 0.02% of the differentially expressed genes).

Enrichment of gene ontology terms provided functional insights on these differentially expressed genes (Supplemental File 10), and these identified broad categories of biological components and processes that are shared among the nectary types. For example, during the development of floral, bracteal, and circumbracteal nectaries those genes that share the preU-postD and preD-postD temporal expression patterns are enriched for components that are integral plasma membrane proteins. In the bracteal nectary, genes belonging to the preU-postD temporal expression pattern are also enriched for catabolic processes related to lipid and pectin metabolism. The remaining terms lacked informative capacity as they are overly enriched in non-descript annotations, such as “response to stimulus.”

### Expression of carbohydrate metabolism and transmembrane transport genes related to nectar production

Because nectar production is heavily dependent on sugar metabolism (Ren et al., 2007; Solhaug et al., 2019) and sugar transport, the RNA-seq data were annotated with respect to starch and sucrose metabolic pathways and transmembrane transporters, using MapMan (Thimm et al., 2004) and gene ontology terms. The resulting gene list was further filtered, selecting those genes that are upregulated in the nectary transcriptomes relative to the adjoining control non-nectary transcriptomes. A secondary filter was also applied to select those genes that display developmental stage-dependent differential expression within a specific nectary. Figure 12 illustrates as a heat map, the temporal expression patterns of the 20 selected genes relative to the secretory stage among the four different nectaries (Supplemental File 11). The sequential order of these genes in Figure 12 is in order of their functionality in the eccrine-based model of nectar secretion [reviewed by (Roy et al., 2017)].

**FIGURE 12.**
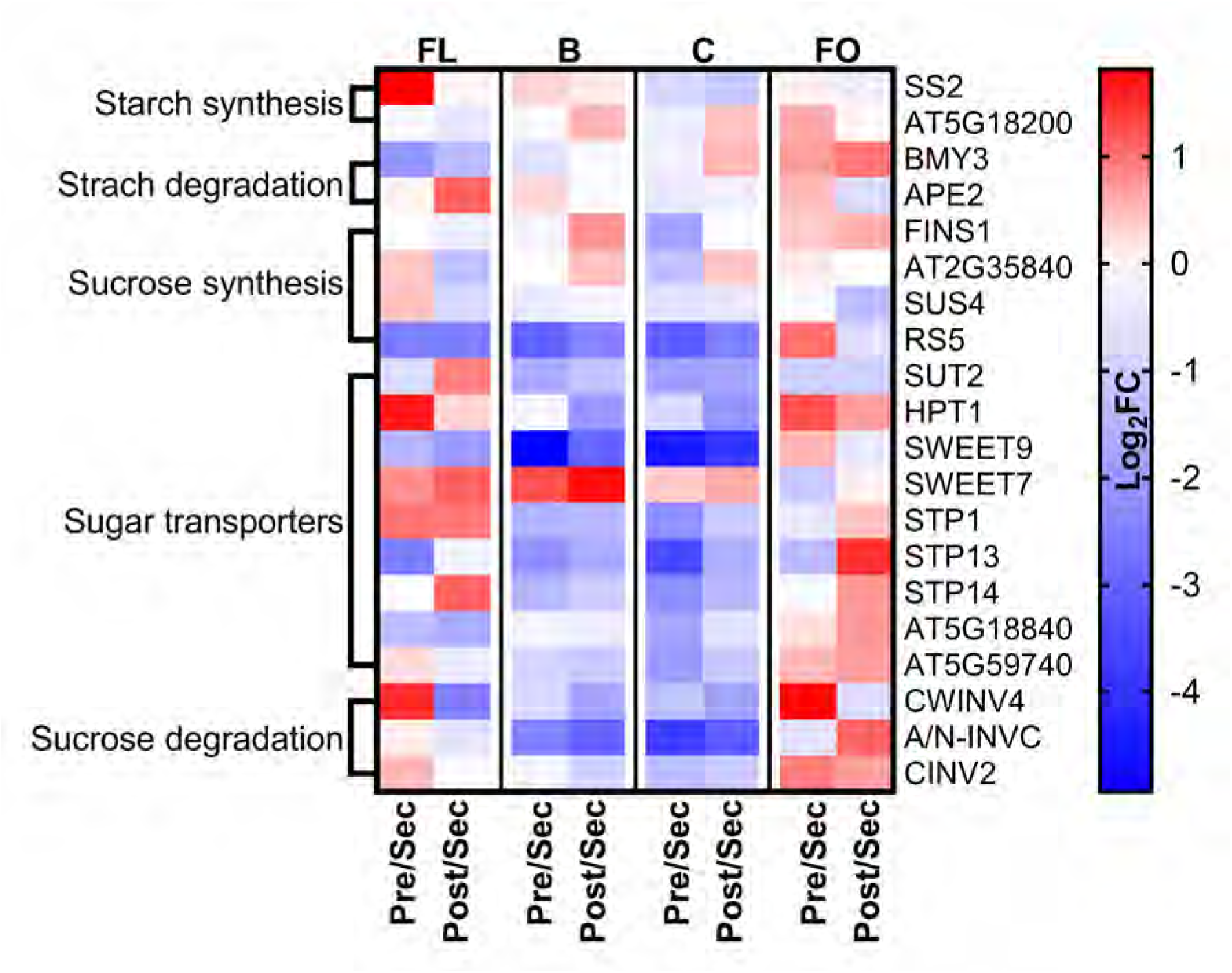
Expression analysis of genes involved in starch and sucrose metabolism. Normalized RNA-seq data was used to generate heat maps of changes in gene expression as each nectary-type transition from pre-secretory to secretory and from secretory to post-secretory stages of development. The blue-red color scale indicates the relative fold-change (FC) between these developmental transitions, on a logarithmic (base-2) scale. Full names for the abbreviations of individual genes are provided in Supplemental File 10. Abbreviations: FL = floral; B = bracteal; C = circumbracteal; FO = foliar; Pre = pre-secretory; Sec = secretory; Post = post-secretory

Consistent with the metabolic events predicted by the eccrine model of nectar production, the floral nectary displays gene expression patterns starting with the upregulation of *SS2* (*Starch Synthase 2*) at the pre-secretory stage, followed by the higher expression of *BMY3*, *SUS4*, *SWEET9*, and *CWINV4* during the secretory stage. In the bracteal and circumbracteal nectaries the expression profiles of these genes deviate from the floral nectary profile. Namely, *BMY3* expression is relatively constant through development, whereas both *SUS4* and *RS5*, involved in sucrose synthesis, are highly expressed during the secretory stage, along with the sugar:proton symporters, *SUT2*, *STP1*, *STP13*, and *STP14*, and a UDP-galactose antiporter (AT5G59740). Thus, these sugar transporters may contribute to the export of sugars during nectar secretion, in addition to *SWEET9*. The expression patterns of these genes in the foliar nectary do not align with the expectation of the eccrine-based model; the exception being the peak expression by *SS2* and a putative galactose-1-phosphate uridyltransferase (AT5G18200) during the pre-secretory stage of development.

Being primarily secretory organs, the nectary transcriptomes are enriched in differentially expressed transmembrane transporters. These include 79 differentially expressed transporters that are predicted to transport sugars, amino acids, water, and various ions (borate, phosphate, hydrogen, calcium, chloride, iron, potassium, and zinc) (Supplemental File 11). As would be expected, the expression of these transporter-coding genes generally peaks during the secretory stage of nectary development (27% of foliar, 39% of floral, 81% of bracteal, and 86% of circumbracteal; Supplemental Fig. 5). Transporters that commonly show peak expression at either the pre-secretory or secretory stage of all four nectary types include those needed for the movement of water via plasma membrane intrinsic proteins (AT2G37170, AT3G53420, AT2G45960). In contrast, the amino acid transporters, *PROT1* (AT2G39890), AT1G47670, and AT3G56200, show temporal differential expression during the development of floral, bracteal and circumbracteal nectaries, but not in the foliar nectary (Supplemental File 11 and Supplemental Fig. 5).

### Upregulation of nitrogen assimilation and amino acid biosynthesis within nectaries

Analysis of the transcriptome data indicate that the floral, bracteal and circumbracteal nectaries display upregulated expression of genes associated with nitrogen assimilation during the secretory stage of development. These genes encode functionalities associated with nitrate transport to the nectary tissue (nitrate transporter *NRT1.5*, AT1G69850), reduction of nitrate to ammonium (nitrate reductase *NR2*, AT1G37130 and nitrite reductase *NIR1*, AT2G15620), and the fixation of ammonium to glutamate by a glutamine synthase (*GLN1*, AT5G37600) (Fig. 12; Supplemental File 11). In foliar nectaries not all of these genes are upregulated, and those whose expression is modulated (for the transport of ammonium and nitrate, by *TIP2;1* (AT3G16240) and *NRT1.2* (AT1G69850), respectively), peak expression occurs during the secretory stage of development.

With amino acids being the second most abundant class of nectar metabolites we examined the nectary transcriptomes for genes associated with amino acid biosynthesis, using MapMan (Thimm et al., 2004) and AraCyc (Mueller et al., 2003). Using the filtering criteria described for the transmembrane transporters, we identified a set of gene products that use glutamate as a substrate for the biosynthesis of amino acids. These genes that function primarily in the biosynthesis of alanine, aspartate, glycine, and branched chain amino acids, show peak expression during the secretory stage of the floral, bracteal and circumbracteal nectaries; aspartate being the prominent amino acid in these nectars. (Fig. 13).

**FIGURE 13.**
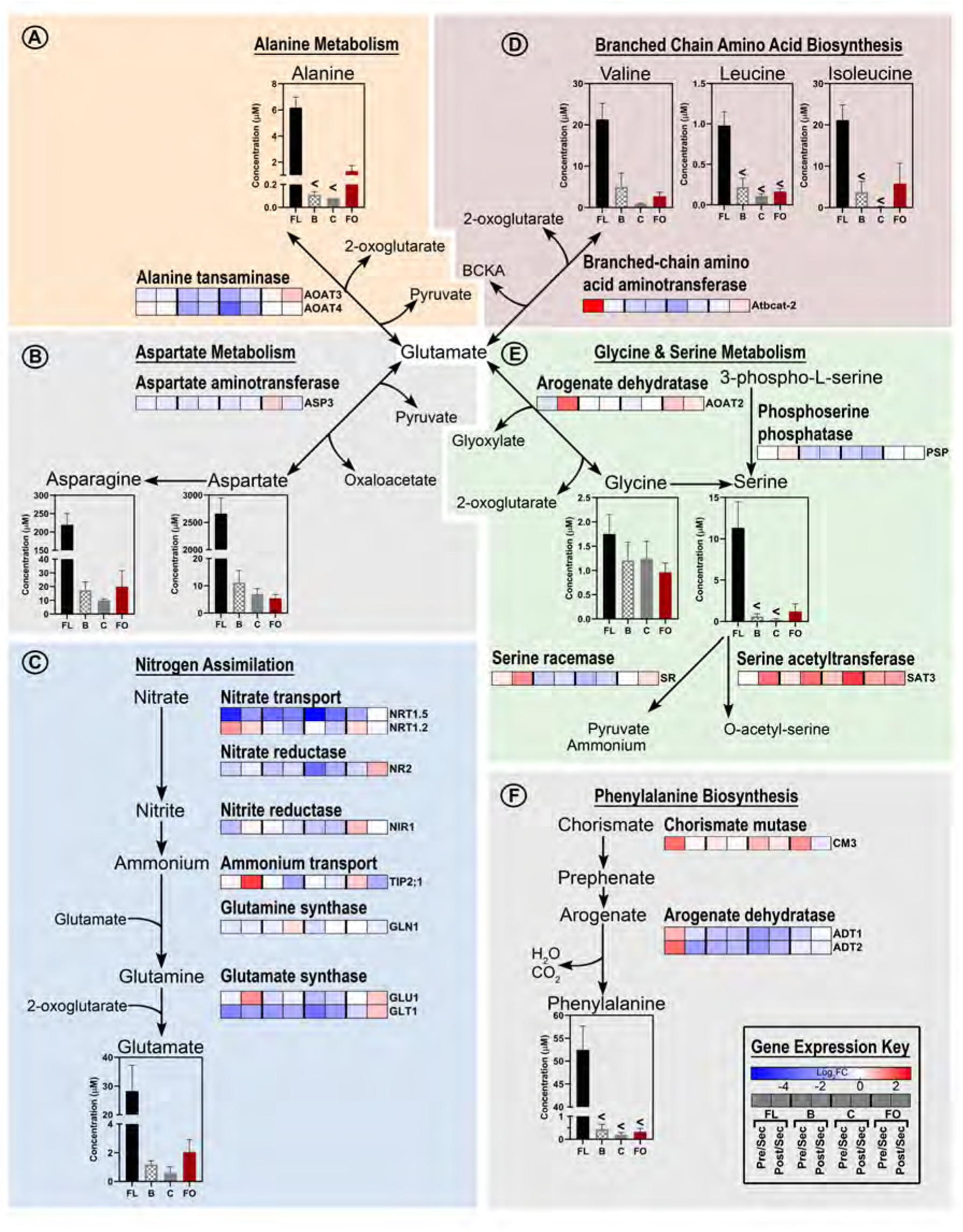
Integration of metabolomics and transcriptomics data to decipher the metabolic processes that support nitrogen assimilation and amino acid biosynthesis in nectaries. Each metabolic module (A-F) integrates metabolomics data of metabolic intermediates and gene expression data of enzymes catalyzing key metabolic processes. The “gene expression key” indicates the logarithmic (base-2) fold-change (Log2FC) between the four nectary types as modulated by developmental transitions. Gene descriptions are provided in Supplemental File 10. Data-bars labeled with the “<” symbol indicate metabolite levels that are below the detection limit of the analytical method. Abbreviations: FL = floral; B = bracteal; C = circumbracteal; FO = foliar; Pre = pre-secretory; Sec = secretory; Post = post-secretory.

### Cell wall and lipid metabolism during nectar secretion

As indicated by the morphological studies of the nectary papillae, we anticipated that genes associated with cell wall and cuticle deposition may show altered expression during development. Such genes were selected based on the spatial and temporal differential expression patterns as revealed by the RNA-seq data, and they were mapped to metabolic networks using MapMan (Thimm et al., 2004). Consistent with expectations, these analyses indicate that during bracteal and circumbracteal nectary development cell wall and cuticle associated genes display temporal differential expression, but this is not the case for floral and foliar nectaries (Supplemental Figs. 6 & 7; Supplemental File 11). Specifically, in both bracteal and circumbracteal nectaries eight genes related to cell wall re-structuring showed statistically significant upregulation during the secretory stage; these include an expansin (*EXLA1*, AT3G45970), and genes required for the synthesis of cell wall components such as callose (*GSL10*, AT3G07160), hemicellulose (*GALT6*, AT5G62620), and pectins (*PME17*, AT2G45220). Likewise, 17 genes related to cuticular lipid metabolism, including fatty acyl elongation, transport of lipids, including cutin (*ABCG11*, AT1G17840) are commonly upregulated in these two nectary-types.

## Discussion

This study represents the first system-based comparison of the four nectary types of *G. hirsutum*. Specifically, we compared and contrasted the nectary morphologies, nectary transcriptomes, and nectar metabolomes of the floral, bracteal, circumbracteal, and foliar nectaries of cotton. These data build upon genetic models for nectar production developed primarily using floral nectaries of Arabidopsis and *Nicotiana* spp., which are nectary tissues containing modified stomata, referred to as ‘nectarostomata’ (Bender et al., 2012, 2013, Carter et al., 1999, 2006, 2007, Carter and Thornburg, 2000, 2004; Hampton et al., 2010; Kram and Carter, 2009; Lin et al., 2014; Liu and Thornburg, 2012; Ren et al., 2007; Ruhlmann et al., 2010; Thomas et al., 2017; Thornburg et al., 2003; Wiesen et al., 2016). Thus, this study evaluates the applicability of the nectar production model developed from studies of floral nectaries to extrafloral nectaries, and nectaries that are composed of secretory trichomes (papillae). The study revealed metabolic processes that are temporally regulated as these papillae nectaries progress from pre-secretion to secretion to post-secretion stages of development. Additionally, regulation of these metabolic processes varies among the three cotton nectary categories, floral, reproductive extrafloral, and vegetative extrafloral. Each of these nectaries have distinct patterns of nectar secretion, with the floral and reproductive extrafloral nectaries following a fixed ontogenetic pattern of secretion and the vegetative extrafloral nectary displaying low constitutive secretion, which is induced upon herbivory (Wäckers and Bonifay, 2004).

### Morphology and ultrastructure of cotton nectaries

Our studies expand upon earlier descriptions of the morphology and ultrastructure of the cotton foliar nectaries (Eleftheriou and Hall, 1983a; Wergin et al., 1975), and extends such studies to the floral nectary and the reproductive extrafloral nectaries (i.e., bracteal and circumbracteal). The four nectaries of *G. hirsutum* share the basic structural components of similar trichomatic nectaries reported in other taxa (*Abutilon* - Kronestedt et al., 1986; *Hibiscus* - Sawidis et al., 1987; *Platanthera* - Stpiczyńska et al., 2005; *Utricularia* - Plachno et al., 2018). Specifically, subnectariferous parenchyma is associated with vasculature. The nectariferous parenchyma is composed of small isodiametric cells with densely-staining cytoplasm, and closely packed papillae glands, which are composed of a single basal cell, variable number of stalk cells, and terminal head cells protruding from the epidermis (Bernardello et al., 2007; Eleftheriou and Hall, 1983a; Fahn, 1979; Findlay et al., 1971b; Kronestedt et al., 1986; Lattar et al., 2018; Sawidis et al., 1987; Wergin et al., 1975). Large phenolic ‘dense’ bodies and calcium oxalate crystals form in all four nectaries and based on their postulated functionality in nectaries of other plant taxa, such as *Glycine*, *Linaria*, *Epipactis*, *Heliocarpus*, *Luehea*, *Ekebergia*, and *Anacardium*, they may confer protection from herbivory (Horner et al., 2003; Jachuła et al., 2018; Kowalkowska et al., 2018; Lattar et al., 2018; Tilney et al., 2018; Tölke et al., 2018).

The ultrastructure of nectariferous tissues reveals abundant ribosomes and a normal complement of organelles, including prominent rough endoplasmic reticulum, abundant mitochondria, scarce plastids, and few Golgi bodies. The abundance of rough endoplasmic reticulum positioned parallel to the cell walls may contribute to vesicle trafficking between cells in support of the granulocrine model of nectar secretion (Eleftheriou and Hall, 1983a). Pit fields of plasmodesmata traverse the cell walls of the nectariferous parenchyma and the inner anticlinal and peridermal walls of the papillae. This distribution of plasmodesmata supports symplastic flow of pre-nectar metabolites, such as sugars, from the associated vasculature to ultimate secretion of nectar from the papillae head cells (Eleftheriou and Hall, 1983a; Findlay et al., 1971a; Wergin et al., 1975). In contrast to the previous studies of foliar nectaries, cell wall ingrowths were observed during the secretory stage of bracteal and circumbracteal nectaries on the distal cell wall of the papillae head cells. These ingrowths in the region of nectar secretion, may facilitate nectar secretion by increasing the surface area (Fahn, 1979; Plachno et al., 2018), which may be particularly important for the reproductive extrafloral nectaries that produce the largest volume of nectar and are active for the duration of fruit maturation (Wäckers and Bonifay, 2004).

During active nectar secretion, the cuticle of the papillae head cells separates from the cell wall and the newly formed subcuticular space, which fills with nectar; microchannels or fractures develop in the cuticle to facilitate the release of nectar from the nectary. This phenomenon commonly occurs in the trichomatic nectaries of a variety of other species, included within Malvaceae (Findlay et al., 1971b; Haratym and Weryszko-Chmielewska, 2017; Kowalkowska et al., 2018; Kronestedt et al., 1986; Lattar et al., 2018; Plachno et al., 2018; Sawidis et al., 1987). Based on previous observations of *Abutilon hydridum* floral nectary papillae, the cuticular channels may function as valves, releasing discrete droplets of nectar once hydrostatic pressure exceeds a threshold (Findlay et al., 1971b).

### Nectar metabolomes reflect the feeding preferences of target facultative mutualists

The distinct nectars produced by *G. hirsutum* floral and extrafloral nectaries parallel the feeding preferences of the pollinating mutualists visiting the floral nectary (honey bees) and the protective mutualists visiting extrafloral nectaries (ants). This variation between floral and extrafloral nectars has previously been reported for a number of species that produce both nectar types on a single plant (Baker et al., 1978). Furthermore, our finding of a unique metabolite profile of floral nectar agrees with prior studies (Butler et al., 1972; Gilliam et al., 1981; Hanny and Elmore, 1974). Specifically, reflecting the known feeding preference of bees (Baker and Baker, 1983; Waller, 1972), which visit cotton flowers, the floral nectar is the most hexose-rich cotton nectar type, containing minimal sucrose, and has the highest abundance and widest variety of amino acids. We also identified GABA as a non-proteinaceous amino acid unique to floral nectar. Based on the fact that phenylalanine and GABA are known to elicit a strong phagostimulatory response in bees, the presence of these floral nectar-specific amino acids may function to attract this pollinator (Hendriksma et al., 2014; Nepi, 2014; Petanidou et al., 2006). GABA may also confer health benefits for bees as GABA-enriched artificial nectar has been shown to increase the locomotion and survival time of bees (Bogo et al., 2019). Leucine and tryptophan may also provide a desirable flavor due to stimulation of sugar chemosensory cells (Shiraishi and Kuwabara, 1970). Lastly, proline was the fourth most abundant amino acid of floral nectar, which is particularly important for bees by providing a rapid energy source for initial flight take-off (Carter et al., 2006; Teulier et al., 2016).

The extrafloral nectars, which function as a reward for mutualist ants, are characterized by higher sucrose content, and a broader distribution of amino acids, which are at lower abundance levels than in floral nectar. These characteristics reflect ant feeding preferences for carbohydrate sources rich in fructose and glucose to sustain worker ant metabolism, which also contain complex mixtures of amino acids, proposed to provide flavor and dietary nitrogen (Blüthgen and Fiedler, 2004; Dussutour and Simpson, 2009; Lanza, 1988). Similar to the floral nectar, extrafloral nectars also contain a high proportion of proline relative to the other amino acids, albeit at a concentration ten-fold less than the floral nectar; proline accounts for 9% to 12% of the amino acids of extrafloral nectar (the fourth most abundant), and accounting for 22% of the amino acids of vegetative extrafloral nectar (the second most abundant in this nectar). The biological effects of proline on ants remains unexplored, but a survey of ant food sources identified proline as the most abundant amino acid (Blüthgen et al., 2004). A final feature which separates extrafloral nectars from floral nectar is the high proportion of non-proteinaceous amino acids largely composed of β-alanine (i.e. 6% to 20% in extrafloral nectar, compared to 0.05% of floral nectar). In addition to the sugar and amino acids, these nectars also contain lipids (Stone et al., 1985), but their role in attraction of mutualists is not clear.

### Nectary capacity for *de novo* amino acid synthesis and transport

As evidenced by the upregulation of the core genes required for nitrogen assimilation, nectaries of cotton, particularly the floral and reproductive extrafloral nectaries, exhibit the capacity to reduce nitrate and incorporate ammonium into organic forms. Specific genes associated with these processes include: nitrate transporters (*NRT1.5, NRT1.2*), nitrate reductase (*NR2*), nitrite reductase (*NIR1*), ammonium transporter (*TIP2-1*), glutamine synthase *(GLN1*), and glutamate synthase (*GLT1, GLU1*) [reviewed by (Dechorgnat et al., 2010)]. While nitrogen assimilation commonly occurs in roots, it also occurs in shoots where photosynthesis can provide energy (Lin et al., 2008; Meyer and Stitt, 2001), but there are no previous reports of these processes occurring in nectaries.

Based on these transcriptomic profiles, one can surmise that nitrate is initially transported through the xylem to the subnectariferous parenchyma by the cotton orthologs of the proton-coupled nitrate transporter *NRT1.5* (Lin et al., 2008; Tsay et al., 2007). Once in the subnectariferous parenchyma, nitrate undergoes two successive reductions to produce ammonium, which is used to assemble glutamine by glutamine synthase (*GLN1*). Glutamate synthase (*GLU1* and *GLT1*) catalyzes the reaction of glutamine with 2-oxoglutarate to form glutamate [reviewed by (Bernard and Habash, 2009)]. Ammonium flux maybe modulated by the tonoplast localized ammonium uniporter *TIP2;1*, a gene upregulated in secretory reproductive extrafloral nectaries (Loque et al., 2005).

Nectary transcriptome data also revealed an upregulation of genes associated with amino acid biosynthesis and amino acid transporters, which are up-regulated at the pre-secretory and secretory stages of nectary development. These changes in expression are consistent with the amino acid profiles of the secreted nectars. For example, expression of *aspartate aminotransferase 3* (*ASP3*, AT5G11520) was highest among all nectaries, at the pre-secretory and secretory stages. This enzyme utilizes the glutamate, produced via ammonium assimilation, to convert oxaloacetate to aspartic acid, one of the most abundant amino acids of floral and extrafloral nectars.

Other such correlations between nectar amino acids and biosynthetic enzymes include phenylalanine and the biosynthetic enzyme, arogenate dehydratase 2 (*ADT2*, AT3G07630), and proline and the proline transporter *PROT1* (AT2G39890) (Yamada et al., 2011). Collectively therefore, these data suggest that cotton nectaries actively assimilate inorganic nitrogen into the amide moiety of glutamine, which functions as the amino group donor for synthesis of additional amino acids such as alanine, glycine, and branched chain amino acids. These amino acids can then undergo symplastic transport to the head cells of the papillae through the action of the upregulated amino acid transmembrane transporters culminating in deposition into the secreted nectars.

### Mechanisms of cotton nectar secretion supported by the transcriptome and papillae ultrastructure

Multiple mechanistic models have been proposed for the biosynthesis of nectar components and release from trichomatic nectaries (Findlay et al., 1971b; Kronestedt et al., 1986; Paiva, 2016). These mechanisms must explain how the nectar components cross the barriers posed by the plasma membrane, cell wall, and cuticle. The potential complexity of this process is multiplied when considering the variation between floral and extrafloral nectaries which have contrasting patterns of nectar secretion and origins of pre-nectar metabolites (i.e. starch storage or lack thereof). The merocrine model (also called the granulocrine model) and the eccrine model are the two models that best align with transcriptomes and ultrastructure of the studied cotton nectaries. These two models likely function in coordination with each other to synthesize nectar components and secrete the metabolites from the nectary tissues. In the merocrine model, nectar metabolites are packaged into vesicles that fuse with the plasma membrane, releasing the nectar components. The eccrine model deviates from the merocrine, in that nectar metabolites are ferried through the plasma membrane by channels and transporters [reviewed by (Roy et al., 2017)]. Currently, the eccrine model is best supported by molecular evidence from floral nectaries of Cucurbitaceae, Brassicaceae and Solanaceae that express five metabolic processes: 1) starch synthesis, 2) starch degradation, 3) sucrose synthesis, 4) export of sucrose into the apoplast via SWEET9, and 5) extracellular hydrolysis of sucrose by CELL WALL INVERTASE4 (CWINV4) (Chatt et al., 2018; Lin et al., 2014; Ruhlmann et al., 2010; Solhaug et al., 2019; Thomas et al., 2017). In both models, prior to the final release of nectar by vesicles or transporters, the pre-nectar metabolites move symplastically through the nectar parenchyma tissues.

The merocrine model is best supported when one considers the ultrastructural analyses of cotton nectaries. Specifically, the prominence of rough endoplasmic reticulum and abundance of vesicles that appear to be fusing to plasma membranes within the nectariferous parenchyma and throughout the papillae are consistent with the importance of vesicle movement to deliver nectar components through the parenchyma cells and out of the nectary papillae. In contrast, the transcriptome expression patterns during the development of floral and reproductive extrafloral nectaries support the eccrine model. Specifically, the expression of genes coding for enzymes and transporters associated with the five metabolic processes that support the biosynthesis and secretion of nectar components. The lack of such an expression pattern during the development of foliar nectaries may be a consequence of the fact that these nectaries produce a steady but low level of nectar, and thus there is no need for a change in a gene expression program that would provide evidence in support of the eccrine model of nectar production.

The upregulation of *SWEET9* and *CWINV4* at the secretory stages of nectary development is also supportive of the eccrine model, and the relative expression levels of these two genes appears to be predictive of whether the nectar product will be hexose-rich. As with Arabidopsis and pennycress nectaries (Bender et al., 2012; Kram et al., 2009; Lin et al., 2014; Ruhlmann et al., 2010; Thomas et al., 2017), which produce hexose-rich nectars, the expression of *SWEET9* and *CWINV4* at the secretory stage of cotton floral nectaries is near equal, and this nectary also produces the most hexose-dominant nectar of cotton. In contrast, the three cotton extrafloral nectaries produce nectars that are more sucrose enriched, and *CWINV4* expression is almost one-sixth the level of *SWEET9* expression. Similarly, such disproportionate expression of *SWEET9* and *CWINV4* has been reported in nectaries of pumpkin, squash, and sunflower, all of which produce sucrose-rich nectars (Chatt et al., 2018; Prasifka et al., 2018; Solhaug et al., 2019).

The eccrine model of nectar deposition has been primarily developed to explain the deposition of the sugar components of nectars. Similarly however, the expression of genes encoding for transporters of the minor components of the nectars would indicate that the eccrine model applies equally to these classes of metabolites. In support of this hypothesis, the transcriptomes of developing cotton nectaries reveals upregulated expression of plasma membrane-H^+^-ATPase, sugar:proton symporters, amino acid transporters, and lipid transmembrane transporters at the secretory stage of nectary development. The expression of such ATPase transmembrane transporters and proton gradients have previously been suggested to facilitate export of nectar metabolites (Bernardello et al., 2007; Chatt et al., 2018; Eleftheriou and Hall, 1983b; Peng et al., 2004; Vassilyev, 2010). Moreover, the occurence of calcium oxalate crystals (druses) around the vasculature and throughout the nectary parenchyma tissues may indicate the need to regulate calcium levels by sequestration as insoluble salts to negate the inhibitory effects of this cation on plasma membrane ATPases (Aguero et al., 2018; Kronestedt et al., 1986; Tölke et al., 2018).

Cotton nectar constituents are ultimately secreted from the papillae head cells, passing through the cell wall and cuticle. Our morphological and anatomical studies of reproductive extrafloral nectaries indicate that this passage is facilitated by microscopic physical alterations in the structure of the cell wall and cuticle. Consistent with the physical alterations of these polymeric structures, the expression of cell wall structural genes is upregulated, which likely contributes to the development of cell wall ingrowths on papillae head cells, increasing the surface area available for the secretion of nectar (Fahn, 1979; Kronestedt et al., 1986; Paiva, 2016). The nectar that has passed through the cell wall appears to accumulate in the subcuticular space between the cell wall and cuticle, generating sufficient hydrostatic pressure to expand this interface, and ultimately be secreted through small pores and fractures in the cuticle. These actions may require the deposition of new cuticular lipids, which may be the driver for the upregulated expression of cuticle deposition genes.

In summary, our combined systems-level studies of the expression of *G. hirsutum* floral and reproductive extrafloral nectaries generated data that support a coordination of merocrine-based and eccrine-based models of nectar synthesis and secretion. The eccrine-based model was primarily developed from studies of eudicot floral nectaries. Therefore, this study has expanded the conservation of the eccrine model, for the first time, to extrafloral nectaries.

## Material and Methods

### Plant materials

Plants were grown in a Conviron Environmental Growth Chambers (0.7 m × 1.8 m × m) that was kept in a cycle of 12 h illumination at 26 °C starting at 6:00 local time, and 12 h darkness at 22 °C. Seeds of *Gossypium hirsutum*, TM-1 were chipped and germinated in 8 cm × 8 cm × 10 cm pots filled with a soil mixture of 3-parts LC8 soil (www.sungro.com) to 1-part sand. Individual seedlings were transplanted into 2-gallon (2A) pots after reaching approximately 30 cm in height, and 10 g of Osmocote Pro 19-5-8 (www.amleo.com) was mixed into the soil mixture per pot. Each growth chamber contained five plants. Plants were watered each day and once per week with a 10% fertilizer solution of Scotts Excel 21-5-20 all-purpose water-soluble fertilizer and Scotts Excel 15-5-15 Cal-Mag water soluble fertilizer (www.jvkbmcmillian).

### Collection of nectary and nectar samples

All nectary and nectar samples were collected from plants after the first flower bloomed, approximately 70 days after sowing. Nectar samples were collected between 10 am and 3 pm local time, using a 5 µL Drummond® Microdispenser (www.drummondsci.com). Nectar samples were first harvested before nectary tissue was excised using a sterile scalpel. Nectary samples were collected from leaves or flowers immediately after removal of each organ from the plants, and the collected nectary tissues were immediately flash-frozen in liquid nitrogen and stored at −80 °C.

In this study, we analyzed four types of nectaries, the floral, bracteal, and circumbracteal nectaries collected from flowers, and foliar nectaries collected from leaves. The developmental trajectory of each nectary type was defined relative to nectar secretion, and are defined as pre-secretory, secretory and post-secretory stages. Thus, in the case of floral nectaries these three developmental stages were collected at 24 h pre-anthesis, at anthesis, and at 24 h post-anthesis. The three equivalent developmental stages for bracteal and circumbracteal nectaries are defined as, a) the “match-head square stage” of cotton square development (Main, 2012), b) anthesis, and c) 19 to 24 days after anthesis. Analogously, the three developmental stages of foliar nectaries were collected from leaves with a midvein length of 5 to 6 cm, a midvein length of 12 to 15 cm, and fully mature leaves that lacked visible nectar deposits.

### Non-targeted metabolomics analysis of nectar metabolites

Two separate GC-MS based methods were employed for non-targeted metabolite profiling of nectar samples. Six replicate nectar samples were collected for each of the four nectar types. Each replicate consisted of pooled nectar, sampled from a minimum of 3 nectaries harvested from two plants on a single day.

One of these analysis methods provided data on the predominant sugars that constitute the nectar (i.e. sucrose, glucose, and fructose). Specifically, 1 µL of nectar, spiked with an internal standard (10 µg ribitol) was dried by lyophilization. The sample was methoximated at 30 °C for 90 min, while continuously shaking with 20 mg mL^-1^ methoxyamine hydrochloride dissolved in pyridine. The methoximated sample was silylated for 30 min at 60 °C with N,O-Bis(trimethylsilyl)trifluoroacetamide and 1% trimethylchlorosilane. Following dilution with mL pyridine, 1-µL sample was analyzed by GC-MS. GC parameters were set to a helium gas flow rate of 1 mL min^-1^, 1 µL injection with a 10:1 split, and a temperature gradient of 100 °C to 180 °C increasing at a rate of 15 °C min ^-1^, then 5 °C min ^-1^ to 305 °C, then 15 °C min ^-1^ to 320 °C, followed by a 5 min hold at 320 °C.

The second analysis method focused on the less abundant constituents of the nectar, which were extracted from a 5-µL aliquot of nectar sample that was spiked with 0.5 µg nonadecanoic acid and 1 µg ribitol, as internal standards. Hot methanol (2.5 mL) was added to the nectar, and the mixture was incubated at 60 °C for 10 min. Following sonication for 10 min at 4 °C, chloroform (2.5 mL) and water (1.5 mL) were sequentially added, and the mixture was vortexed. Centrifugation separated the polar and non-polar fractions, and the entire non-polar fraction and half of the polar fraction was recovered to separate 2 mL screw-cap glass vials and dried by lyophilization. The polar fraction underwent methoximation as previously described, and both polar and non-polar fraction were silylated for 30 min at 60 °C with N,O-Bis(trimethylsilyl)trifluoroacetamide and 1% trimethylchlorosilane.

The derivatized metabolites (the sugars, polar, and non-polar fractions) were analyzed using an Agilent Technologies Model 7890A GC system equipped with an HP-5ms (30 m, 0.25 mm, 0.25 µm) column that was coupled to an Agilent Technologies 7683B series injector and Agilent Technologies Model 5975C inert XL MSD with Triple-Axis Detector mass spectrometer (www.agilent.com). Chromatography parameters for the polar and non-polar fractions were set to a helium gas flow rate of 1 mL min^-1^, 2 µL injection, with a temperature gradient of 80 °C to 320 °C increasing at a rate of 5 °C min ^-1^, followed by a 9 min hold at 320 °C. The polar fractions were analyzed using a “heart-cut” method which diverted gas flow to an FID detector during elution times for fructose, glucose, and sucrose. Deconvolution and integration of resulting spectra were performed with AMDIS (Automated Mass Spectral Deconvolution and Identification System) software (Stein, 1999). Analyte peaks were identified by comparing mass spectra and retention indices to the NIST14 Mass Spectral Library and authentic standards when possible to confirm identification.

### Amino acid analysis

Analysis of amino acids was performed using the Phenomenex EZ:Faast^TM^ kit for free amino acids (www.phenomenex.com). Six replicate samples for each nectar type were collected as described previously. Due to low volume of nectar produced by the foliar nectary, these nectar samples were pooled from a maximum of 90 nectaries, collected from 6 separate plants. Each sample (20 µL nectar per extraction) was subjected to solid phase extraction and derivatized according to the manufacturer’s instructions, with one adjustment: after addition of the norvaline internal standard (5 nmol) to each sample, 125 µL of 10% propanol/20 mM HCl was added to acidify the sample. Following derivatization, samples were concentrated by evaporation under a stream of nitrogen gas before amino acids were analyzed using an Agilent Technologies model 6890 gas chromatograph with a ZB-AAA 10 m × 0.25 mm amino acid analysis column coupled to a model 5973 mass selective detector capable of electrical ionization (EI). The GC-MS instrument settings followed the manufacturer’s recommendations.

### Statistical analysis of cotton nectar metabolites

For each metabolite, the natural logarithm of normalized metabolite level was averaged over the six replicates for each nectar type. Separately for each metabolite, a linear model with one mean per species and constant error variance was fitted to the metabolite response values. As part of each linear model analysis, F-tests for contrasts among the 4 nectar type means were conducted to identify differences in average response between each pair of nectar types. The 197 p-values for each comparison (one p-value per metabolite) were adjusted to obtain approximate control of the false discovery rate at the 0.05 level (Benjamini and Hochberg, 1995).

Similarities and differences among metabolites between different nectary types were visualized by pair-wise volcano plot comparisons and hierarchical agglomerative clustering. To perform clustering, the estimated nectar type response means were first standardized within each metabolite to obtain a standardized response profile across nectar types for each metabolite. Then dissimilarity between each pair of metabolites was computed as the Euclidean distance between the standardized response profiles. Clustering based on these pairwise dissimilarities places two metabolites in the same cluster if their estimated nectar type response means are highly correlated across sections. Although hierarchical clustering groups the metabolites into any number of clusters, a total of 16 clusters were selected to display and summarize the results, striking a balance between high within-cluster consistency and low between-cluster similarity.

### Mass spectrometric imaging of nectary metabolites

Nectary tissue was excised from plants and immediately embedded in a 2% solution of carboxymethylcellulose sodium medium viscosity in a disposable base mold (7 × 7 × 5 mm) and flash-frozen with liquid nitrogen. Triplicate samples of all four nectary types (floral, circumbracteal, bracteal, and foliar) at the pre-secretory and secretory stages were similarly prepared. Base molds were allowed to set at −20 °C for about 18 h, before 20 µm transverse cryosections were collected. During sectioning, the embedded tissue blocks were mounted on the cryostat using optimal cutting temperature compound, and sections were collected on 12 mm carbon adhesive tabs (Electron Microscopy Sciences; cat. # 77825-12; www.emsdiasum.com/microscopy/). Sections were dried for 1 h by lyophilization and visually imaged with a Zeiss AxioZoom (www.zeiss.com). Well preserved sections were placed onto indium tin oxide coated glass slides 75 × 25 mm (Bruker, Billerica, MA; cat. #8237001; www.bruker.com). Sections where then coated with a matrix using an oscillating capillary nebulizer sprayer (Hansen and Lee, 2017). The matrix was composed of 4 mL of 5 mg mL^-1^ 1,5-diaminonaphthalene dissolved in acetonitrile, 2 mL methanol, and 2 mL water, and it was applied at a rate of 4 mL h^-1^ in 0.30 mL steps. After matrix application, samples were dried overnight in a desiccator.

MALDI-MS imaging was performed using a Bruker SolariX FT-ICR MS instrument equipped with a 7.0 tesla superconducting magnet. MALDI-MS data was acquired in negative ion mode with a mass range from *m/z* 73 to 1000, collecting 2 megabytes of data points per scan. The laser was set to raster at 25 µm spots, and *flexImaging* software (www.bruker.com) was used to collect and analyze the imaging data. Agilent MassHunter software and the METLIN Metabolomics Database and Library with ppm tolerance set to 8 were used to identify *m/z* values of interest.

### Light microscopy and histochemistry

Pre-secretory and secretory stage nectaries were fixed for several days at 4 °C, in a solution of 3% (w/v) glutaraldehyde and 2% (w/v) paraformaldehyde in 0.1M sodium cacodylate buffer, pH 7.2. Samples were dehydrated in a graded ethanol series (50% - 100%), followed by infiltration and embedding over five days in LR White resin. For replication purposes a minimum of four nectaries per nectary-type where imbedded at each developmental stage. Resin blocks were polymerized at 55 °C for 72 h. Histological sections were cut at 1.3 µm thickness using a Leica UC6 ultramicrotome (www.leica-microsystems.com). Sections were dyed with Toluidine Blue O for general contrast and Periodic Acid Schiff’s (PAS) technique for starch and other water-insoluble carbohydrates (Ruzin, 1999). Digital images were collected using a Zeiss Axiocam HRC camera (www.zeiss.com) on an Olympus BX-40 compound microscope (www.olympus-ims.com) in bright-field mode.

### Transmission electron microscopy

A minimum of four nectaries, of the four nectary types (foliar, bracteal, circumbracteal, and floral), harvested at the secretory stage, were fixed for several days at 4 °C, in a solution of 3% (w/v) glutaraldehyde and 2% (w/v) paraformaldehyde in 0.1 M sodium cacodylate buffer, pH 7.2. Samples were washed with several changes of 0.1 M sodium cacodylate buffer, pH 7.2, and then fixed in 1% osmium tetroxide in 0.1 M sodium cacodylate buffer for 1 h at room temperature. The samples were *en block* stained for 2 h with aqueous 2% uranyl acetate, and then dehydrated in a graded ethanol series (50% - 100%). Following a transition into ultra-pure acetone, and infiltrating, the nectaries were embedded with Spurr’s hard epoxy resin (www.emsdiasum.com). Resin blocks were polymerized for 48 h at 70 °C. Thick sections (1 µm) to check fixation quality and ultrathin (90 nm) sections were made using a Leica UC6 ultramicrotome (www.leica-microsystems.com). Ultrathin sections were collected onto carbon-film, single-slot copper grids and images were captured using a JEM 2100 200kV scanning and transmission electron microscope (www.jeol.com).

### Scanning electron microscopy

A minimum of four nectaries per nectary type and at the pre-secretory and secretory stages of development, were fixed for several days at 4 °C in formalin-acetic acid-alcohol. They were dehydrated in a graded ethanol series (50%, 70, 95, 100, 100 ultra-pure twice). Samples were critical point-dried using a Denton Drying Apparatus, Model DCP-1 (www.dentonvacuum.com). The dried specimens were mounted on aluminum stubs with 12 mm circular carbon adhesive tabs and colloidal silver paint (www.emsdiasum.com). Samples were sputter coated with 30 nm platinum using a Cressington HR208 Sputter Coater (www.cressington.com). Images were captured using a Hitachi SU-4800 field emission SEM at 10 kV (www.hitachi-hightech.com).

### RNA isolation, sequencing, and informatics

Triplicate RNA samples were isolated for each of the nectary types. Each replicate was a pool of approximately 2-4 floral or 10-15 of each of the extrafloral nectaries. Tissue was transferred with clean forceps into a 2 mL Lysing matrix A tube (MP Biomedicals; Ref # 6910-500; www.mpbio.com), resting in a liquid nitrogen bath and containing a ceramic bead. The tubes were quickly transferred to a QuickPrep adaptor (containing dry ice) and attached to the FastPrep 24™-5G (www.mpbio.com) benchtop homogenizer for tissue-pulverization. The samples were subjected to 5-6 pulverization cycles of 40 sec each, at 6 m/sec, with each cycle interjected with a period of immersion in liquid nitrogen and refilling the adaptor with dry ice. Post-pulverization, 600 µL of the RNA lysis buffer of the Quick-RNA™ MiniPrep kit (Zymo Research; Cat# R1054; www.zymoresearch.com) was quickly added to the Lysing matrix tube and the tubes were vortexed. This was followed by the addition of 50 µL of the Plant RNA Isolation Aid (Thermo Fisher Scientific, Cat#AM9690; erstwhile Ambion) to remove common plant contaminants such as polyphenolics and polysaccharides. Quick-RNA™ MiniPrep kit directions were followed for RNA isolation. Agarose gel electrophoresis and UV spectrophotometry were used to assess RNA quality, prior to submission to the University of Minnesota Genomics Center for barcoded cDNA library creation and Illumina HiSeq 2500 sequencing. This produced over 360 million 125-bp paired-end reads with a target insert size of 200 bp and generated ≥24 M reads for each sample, and the average quality scores were above Q30. A few samples did not yield suitable sequencing libraries, and thus were omitted from the analysis.

The reads were mapped to the UTX-JGI *Gossypium hirsutum* genome (v1.1) and predicted transcripts using NCBI’s BLASTN (Camacho et al., 2009). The UTX-JGI annotation was used to map read counts to Arabidopsis genes (Araport 11). Read counts were upper-quartile normalized, and pairwise differential expression tests were performed using a negative binomial distribution with DESeq (Anders and Huber, 2010). The resulting p-values were filtered by restricting to genes with a 50% or greater change in mean normalized counts. The Benjamini-Hochberg method was used to control the false discovery rate at the 0.05 level (Benjamini and Hochberg, 1995).

Differentially expressed genes were identified by filtering the DESeq results within R and categorized (e.g., upregulated during the secretory stage); these categories were visualized by generating Venn diagrams using InteractiVenn (Heberle et al., 2015). Gene Ontology (GO) enrichment analysis of the nectary transcriptome was implemented using topGO: Enrichment Analysis for Gene Ontology (Alexa and Rahnenfuhrer, 2016) with prior gene-to-GO term mapping completed using GO.db (Carlson, 2016). A Fisher’s exact test was completed to test for enrichment of GO terms in specific expression pattern groups, using the complete set of 16,958 Arabidopsis orthologs as the baseline for this comparison.

Mapping genes to metabolic pathways used MapMan (Thimm et al., 2004) with the base pathways and mappings files for Arabidopsis. Hierarchical clustering based on one minus Pearson correlation of the log_2_ normalized read count of selected metabolic pathways or functionalities was completed using Morpheus (https://software.broadinstitute.org/morpheus).

### Quantitative Real Time PCR Validation

The same RNA samples used for RNA-seq analyses were subjected to cDNA preparation using the BioRad iScript cDNA synthesis kit (Catalog # 1708890), with 1 μg of RNA used for cDNA preparation. Expression patterns for representative genes that displayed stage specific variation via RNAseq analyses were validated by quantitative RT-PCR using Agilent Brilliant III Ultra-fast SYBR Green QPCR Master Mix (Catalog #600882) and a final cDNA template concentration of 2ng/μl. Expression values are expressed as fold-change relative to the presecretory stage and are based on the delta delta Ct values obtained from the normalized Ct values for each gene. Gene expression was normalized to a gene encoding a 40S ribosomal protein S3-2-like gene (Cotton gene ID= Gohir.D05G034300.1, 1). This gene was chosen as the internal reference based on its stable expression level in floral and bracteal nectary samples across stages in our RNA-seq dataset. Primer sequences for each gene are provided in Supplemental File 12.

### Data availability

Raw sequence reads are available at the National Center for Biotechnology Information Sequence Read Archive under GEO accession number GSE113373. Metabolomics data is publicly available in the PMR database (http://metnetweb.gdcb.iastate.edu/PMR/).

## Acknowledgements

We thank Drs. Ann Perera, Lucas Showman, and Kirthi Narayanaswamy of the W.M. Keck Metabolomics Research Laboratory, Iowa State University for technical support in metabolomics analyses. We are grateful to Tracey P. Stewart and Randall Den Adel, of the Roy J. Carver High Resolution Microscopy Facility, Iowa State University for technical support in microscopic analyses. We also thank Dr. Daniel S. Nettleton and Xingche Guo for conducting hierarchical cluster analysis of metabolomics data and Anthony Schmitt for assistance with RNA preparation. This work was supported by the National Science Foundation award #IOS 1339246 to BJN and CJC.

